# Transposon Regulation in the *Caenorhabditis elegans* Germline and Soma

**DOI:** 10.1101/2025.10.09.681459

**Authors:** Cindy Chang, Dong Cao, Daniel J. Pagano, Scott Kennedy

## Abstract

Transposons are parasitic nucleic acids present in most genomes. The ability of transposons to mobilize makes them a source of genetic diversity and a threat to genome integrity. Interestingly, mutations in the *C. elegans* gene *mut-2/rde-3* increases the rate of transposition in the germline, but not in the soma, suggesting that the *C. elegans* germline and soma employ different strategies to regulate *Tc1* transposition. Here, we develop fluorescence reporters for studying DNA transposon regulation in living *C. elegans* in a tissue-specific manner and we use candidate gene testing and genetic screening approaches to identify factors that regulate *Tc1* mobility in the germline and/or the soma of the animal. We find that both cytoplasmic and nuclear components of the RNA interference (RNAi) pathway silence *Tc1* in the germline, but not in the soma. We identify a novel pathway involving the *C. elegans* ortholog of HNRNPC, and a gene we term suppressor of transposon mobilization (*stm*)*-1*, which regulates *Tc1* primarily in the soma, likely by binding *Tc1* RNA and preventing its splicing. Our findings reveal tissue-specific strategies for regulating parasitic nucleic acids and pave the way for future studies exploring how and why different tissues adopt different transposon silencing systems.

## Introduction

Transposons are mobile genetic elements present in almost all eukaryotic genomes, where they often represent a large fraction of the genome (3-85%) [1]. For example, 40-50% of the human genome is composed of active transposons or inactive transposon relics [1]. Transposon insertions can disrupt host-gene functions, and high-copy transposons make genomes vulnerable to chromosome rearrangements [2]. To limit these deleterious effects of transposons, organisms have evolved defense mechanisms, including DNA, chromatin, and small RNA-based systems, which collectively silence transposon expression and restrict transposon mobility [3–6].

Transposons can be divided into two classes; The retrotransposons and the DNA transposons. Retrotransposons use an RNA-intermediate, which is subjected to reverse transcription, prior to transposition. Retrotransposons replicate via an RNA intermediate and are the most abundant type of transposon in the human genome, where they comprise >40% of the genome [7]. DNA transposons employ a copy-and-paste mechanism for transposition. DNA transposons are inactive in the human genome, yet relics of ancient DNA transposons comprise (∼2-3%) of the genome [7]. In the *C. elegans* genome, DNA transposons are the more common transposon, comprising 12% of the genome [8,9]. The most active and abundant of these elements are the *Tc1/mariner* class DNA transposons [9]. *Tc1/mariner* DNA transposons typically produce a single RNA that, after splicing and polyadenylation, is translated into transposase, which encodes the enzyme responsible for transposon excision and re-insertion [10,11].

The evolutionary impacts of transposition in the germline and soma differ for transposons and their hosts. For transposons, germline transposition is necessary for survival, while somatic transposition is likely of little to no benefit as it does not lead to a heritable change in copy number [12]. For hosts, the short-term fitness costs associated with transposition in the germline or soma are typically negative because transposition is more likely to disrupt, rather than improve, cell functions. Indeed, transposition in both the germline and soma have been linked to disease in humans [13–16]. And the impacts of transposition in the germline are likely to be more severe for the host than transposition in the soma because germline transposition impacts all cells (germ and soma) and these impacts are heritable. The differing goals of transposons and their host cells, as well as the differing impacts of transposition in the soma or the germline, suggest that the strategies employed by multicellular organisms to regulate their transposons may differ in the germline and soma. Consistent with this idea, in the *Drosophila* soma and germline, DNA transposons termed *P* elements are regulated by distinct alternative splicing programs [17]. Additionally, the Gypsy retrotransposon is inhibited by the piRNA *flamenco* locus in somatic cells of the germline but not in germ cells themselves [18]. In *C. elegans*, the rate at which the *Mariner* class DNA transposon *Tc1* transposes is thought to be 1000x higher in the soma than in the germline, and loss of the RNA interference (RNAi) factor RDE-3/MUT-2 increases the rate of *Tc1* transposition in the germline but not in the soma [19–21]. These observations hint that, as in flies, distinct mechanisms may exist to regulate transposons in the germline and soma of *C. elegans* [21].

Heterogenous nuclear ribonucleoprotein C (HNRNPC) is an abundant, nuclear protein that associates with most pre-mRNA [22,23]. HNRNPC regulates many aspects of gene expression, including pre-mRNA splicing as well as cleavage and polyadenylation [23–25]. Transposons often harbor cryptic splice sites, which can lead to aberrant splicing of transposon sequences into host mRNAs, a process referred to as transposon exonization. In mammals, HNRNPC prevents the exonization of *Alu* retrotransposons, which are the most abundant retrotransposon in the human genome [26,27]. Current models posit that HNRNPC binds uridine-rich tracts in *Alu* RNA [23,27–33] to prevent the splicing factor U2AF65 from binding these elements, thus inhibiting *Alu* exonization [27]. Whether the role of HNRNPC in transposon regulation is conserved or whether the pathway is differentially employed in soma/germline contexts is not yet known.

Here we report the development of reporter genes for visualizing *Tc1* excision in the soma and germline of living *C. elegans*. We use these reporters to identify cytoplasmic and nuclear RNA interference (RNAi) factors that limit *Tc1* excision in the germline, but not in the soma. Additionally, we use a genetic screening approach to identify two nuclear-localized proteins, including the *C. elegans* ortholog of HNRNPC, HRPC-1, which we show are important for repressing *Tc1* in the soma, likely by binding and preventing the splicing of *Tc1* RNA. Thus, the *C. elegans* germline and soma employ distinct strategies to control their transposons; RNAi in the germline; and an HNRNPC-based pathway in the soma.

## Results

### Reporter genes to visualize transposon excision in *C. elegans*

*C. elegans Tc1* is a *Mariner* class DNA transposon which is mobile in the *C. elegans* soma and germline [10]. To explore the biology of transposons in an intact animal system, we engineered a reporter gene under the control of an ubiquitous promoter (*eft-3*p) to drive expression of mScarlet-I in all cells of the *C. elegans* soma and germline. A *Tc1* element was then inserted 6 nucleotides upstream of the *eft-3*p*::mScarlet-I* start codon (Figure 1A) with the expectation that: 1) the transposon would likely interfere with mScarlet-I expression; 2) transposon excision would permit mScarlet-I expression in those cells in which *Tc1* had excised; and 3) this would allow us to monitor transposon activity in all cells and during all stages of development of a living animal. We used the CRISPR/Cas9 system to insert the *eft-3*p::*Tc1::mScarlet-I* reporter into the *C. elegans* genome. A control mScarlet-I reporter gene lacking a *Tc1* element was inserted into the same chromosomal site. Fluorescence imaging revealed that, as expected; 1) mScarlet-I was expressed in most/all cells in all *eft-3*p*::mScarlet-I* animals; and 2) the addition of the *Tc1* element inhibited this mScarlet-I expression (Figure 1B). In 77% of *eft-3*p::*Tc1::mScarlet-I* animals, no mScarlet-I was observed in any cell of the germline or soma (Figure 1B). In 23% of animals, 1-2 somatic cells expressed mScarlet-I (Figure 1B and see below).

**Figure 1.**
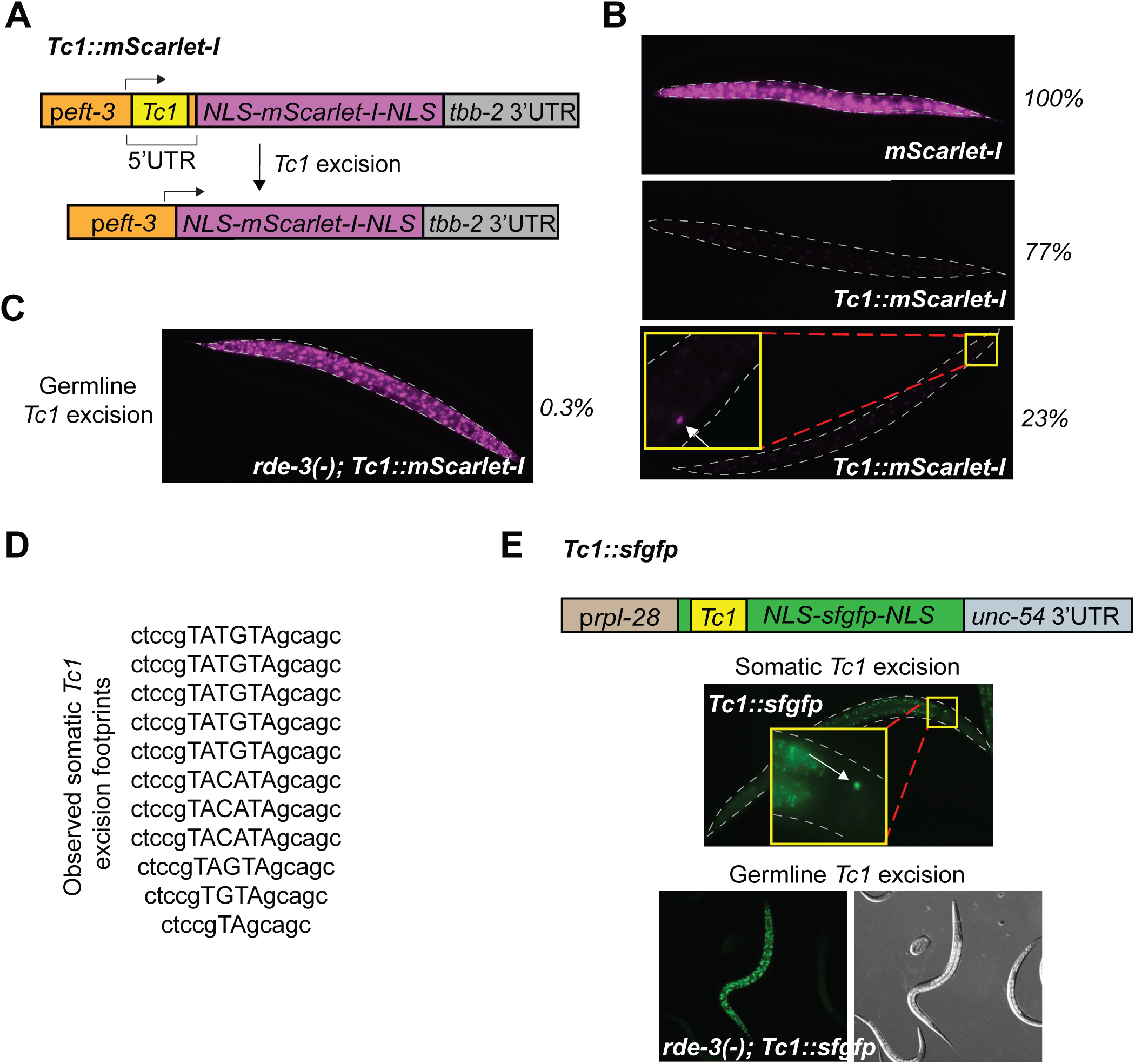
Reporter genes designed to detect *Tc1* excision in living animals. **(A)** Schematic of *Tc1::mScarlet-I* reporter gene engineered to monitor *Tc1* excision. NLS, nuclear localization signal; UTR, untranslated region. Rightward arrow indicates transcription start site and direction of transcription. **(B)** (Top) Fluorescence micrograph of representative *eft-3*p*::mScarlet-I* (*mScarlet-I*) animal. Percentage indicates percentage of *eft-3*p*::mScarlet-I* animals expressing mScarlet-I in all cells. (Middle, Bottom) Fluorescence micrographs of representative *eft-3*p*:: Tc1::mScarlet-I* (*Tc1::mScarlet-I*) animals. 77% of *Tc1::mScarlet-I* animals do not show any expression of mScarlet-I; 23% of *Tc1::mScarlet-I* animals have somatic expression of mScarlet-I. Magnification of mScarlet-I positive cell (arrow) is shown. **(C)** Germline excision of *Tc1* from *Tc1::mScarlet-I* in *rde-3(-)* animal. mScarlet-I expression is observed in all cells of the germline and soma. Percentage indicates the rate of germline excision from *Tc1::mScarlet-I* in *rde-3(-)* animals. During this study, no germline *Tc1* excision was ever observed in wild-type animals. **(D)** Sanger sequencing of *Tc1* excision events from PCR on DNA isolated from *glp-1*; *Tc1::mScarlet-I* animals raised at 25℃, which ablates most germ cells (see Materials and Methods). The PCR strategy allows relatively rare *Tc1* excision events in the soma to be amplified because full-length *Tc1* containing DNA is poorly amplified by the PCR reaction. **(E)** (Top) Schematic of *Tc1::sfgfp* reporter gene engineered to monitor *Tc1* excision. (Middle) Somatic excision from *Tc1::sfgfp* in wild-type animals (arrow, somatic cell expressing sfGFP). (Bottom) Germline excision of *Tc1* from *Tc1::sfgfp* in *rde-3(-)* background.

### *Tc1* reporter genes recapitulate transposon behaviors

We asked if the *Tc1::mScarlet-I* reporter gene recapitulated known properties of *Tc1* and, therefore, could be used to study transposon biology in a living animal. *Tc1* excision in somatic cells would be expected to produce animals expressing fluorescent protein in one or more somatic cells. As outlined above, we observed that 23% of *Tc1::mScarlet-I* animals expressed high levels of mScarlet-I in one or more somatic cells, which suggests that *Tc1* had excised in these cells (Table 1 and Figure 1B). The rate of putative *Tc1* excision in the soma was determined by dividing the total number of animals possessing one or more mScarlet-I positive cells (1304) by the total number of somatic cells scored (≅4.91×10^5^) (Table 1). This analysis set the somatic excision rate of *Tc1::mScarlet-I* at ≥ 2.7 x10^-4^, which is consistent with previous reports for endogenous *Tc1* element activity in the soma (approximately 10^-3^) [19,34]. Four additional lines of evidence show that the *Tc1::mScarlet-I* reporter gene recapitulates known aspects of *Tc1* biology in the *C. elegans* soma and germline.

**Table 1.**
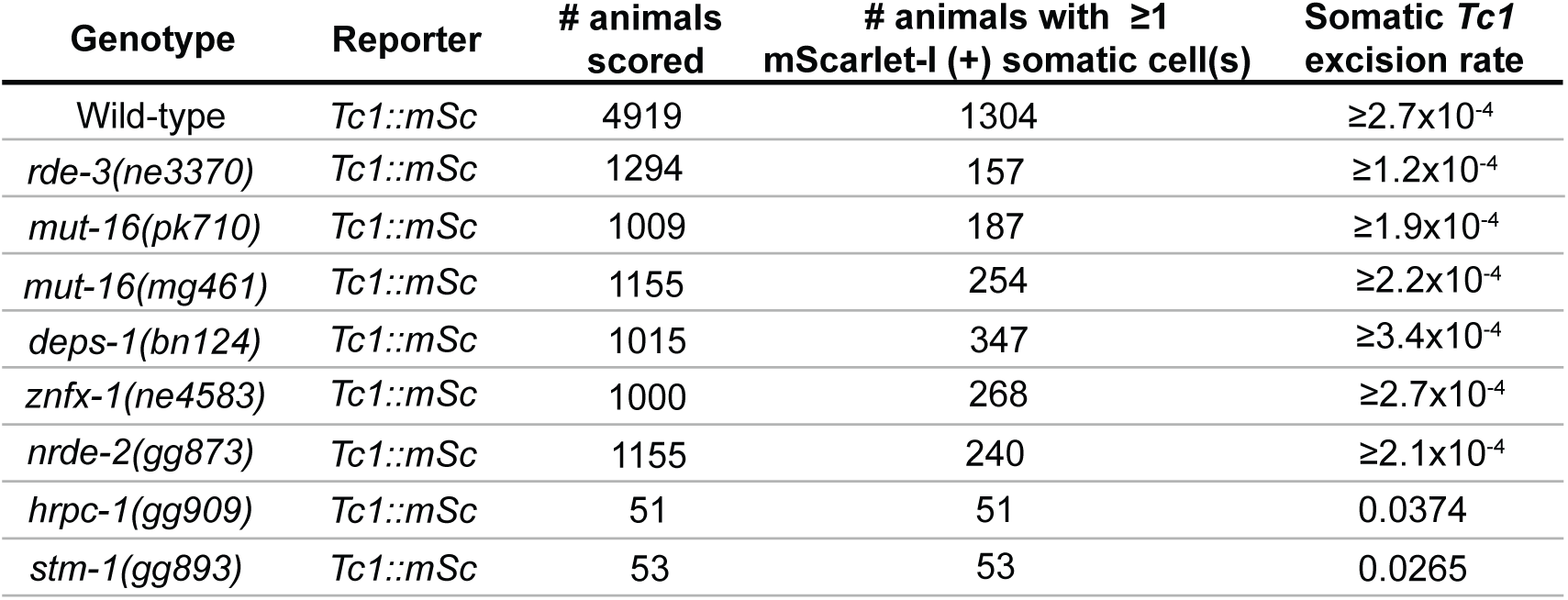
Somatic *Tc1* excision rates. Somatic *Tc1* excision events were quantified by counting the number of somatic cells expressing mScarlet-I (indicated as mScarlet-I (+)) using the *Tc1::mScarlet-I* reporter gene. Somatic *Tc1* excision rates were estimated using the following formula: (# of animals with mScarlet-I(+) cells) x (# of mScarlet-I(+) cells per animal) / ((total # of animals scored) x (1000 = # somatic cells/animal)). For wild-type, if an animal had any mScarlet-I (+) cells the animal typically (73%) had only one mScarlet-I(+) cell. The remaining animals had two (10%) or three (13%) mScarlet-I(+) cells. For these animals, the number of mScarlet-I(+) cells per animal was set as 1 in the above calculation, and the “≥” symbol was used to account for animals that had more than one mScarlet-I(+) cell. For *hrpc-1* and *stm-1* mutants, all animals showed *Tc1* excision and the number of mScarlet-I(+) cells per animal was 37.4 and 26.5 respectively (See Fig 3B). Data are aggregates from at least 3 independent experiments per genotype. *Tc1::mSc*, *Tc1::mScarlet-I*.

First, *Tc1* excision is reported to be ∼1000 fold lower in the *C. elegans* germline than in the soma [19,35,36]. The rate of germline excision for *Tc1* in wild-type *C. elegans* germ cells is reportedly ∼10^-7^ [37]. *Tc1* excision from *Tc1::mScarlet-I* in germ cells would be expected to result in animals that express fluorescent protein in all cells of the germline and soma and this expression state should be heritable. *Tc1::mScarlet-I* germline excision rates were determined by growing animals to high density over several generations and then determining the percentage of plates containing one or more animals expressing mScarlet-I in all cells of the soma and germline, which is indicative of germline transposition. A Poisson distribution method (see Materials and Methods) was used to determine maximum excision rates. Using this approach, we calculated a germline excision rate for *Tc1::mScarlet-I* animals of <2.2×10^-7^ (Table 2), which is similar to the published rate of ∼10^-7^ for endogenous *Tc1* elements [37].

**Table 2.** Germline *Tc1* excision rates. Germline *Tc1* excision events were detected by counting animals in *Tc1::mScarlet-I* or *Tc1::sfgfp* backgrounds (specified in “Reporter used” column) that heritably express mScarlet-I or sfGFP in all cells. Animals were singled onto plates and allowed to grow until plates were populated by F1 or F2/3 broods as indicated. Populations were scored for presence or absence of one or more animals expressing fluorescence in all cells, indicating at least one germline excision event. For genotypes with high germline excision rates (*rde-3(ne3370)*, *mut-16(pk710)*, *deps-1(bn124)*), plates were scored in the F1 generation. For genotypes with low or undetectable germline excision rates (wild-type, *mut-16(mg461)*, *znfx-1(ne4583)*, *nrde-2(gg873), hrpc-1(gg909), stm-1(gg893)*), plates were scored in the F2/3 generations. *Tc1::sfgfp* animals were all scored in F2/3 generations as the *Tc1* excision excision rate from this reporter is lower than from *Tc1::mScarlet-I.* The proportion of plates for each genotype showing germline excision event(s) and the average number of animals per plate at the time of scoring for each genotype were used to calculate the germline excision rate using a Poisson distribution method (see Materials and Methods for formula explanation). For genetic backgrounds where the rate of germline excision was so low that no excision events were ever observed, the Poisson distribution method was used to set a maximum rate of germline excision (indicated with “<”). Data are aggregates from at least 3 independent experiments per genotype. ND, not determined. *Tc1::mSc*, *Tc1::mScarlet-I*. *gg1160* and *gg1161* are independent alleles created by CRISPR/Cas9, both equivalent to the *ne298* allele [42,61].

| Genotype | Reporter | # plates scored | # animals per plate (F1) | # animals per plate (F2/3) | # plates with animal(s) expressing mSc/sfGFP in all cells | Germline *Tc1* excision rate |
| --- | --- | --- | --- | --- | --- | --- |
| Wild-type | *Tc1::mSc* | 167 | ND | 27140 | 0/167 | $<2.2 \times 10^{-7}$ |
| *rde-3(ne3370)* | *Tc1::mSc* | 81 | 322 | ND | 54/81 | $3.4 \times 10^{-3}$ |
| *mut-16(pk710)* | *Tc1::mSc* | 32 | 416 | ND | 19/32 | $2.2 \times 10^{-3}$ |
| *mut-16(mg461)* | *Tc1::mSc* | 34 | ND | 29958 | 0/34 | $<1 \times 10^{-6}$ |
| *deps-1(bn124)* | *Tc1::mSc* | 33 | 1183 | ND | 14/33 | $4.7 \times 10^{-4}$ |
| *znfx-1(ne4583)* | *Tc1::mSc* | 36 | ND | 22513 | 10/36 | $1.4 \times 10^{-5}$ |
| *nrde-2(gg873)* | *Tc1::mSc* | 30 | ND | 22026 | 5/30 | $8.3 \times 10^{-6}$ |
| *hrpc-1(gg909)* | *Tc1::mSc* | 61 | ND | 16298 | 0/61 | $<1 \times 10^{-6}$ |
| *stm-1(gg893)* | *Tc1::mSc* | 35 | ND | 23819 | 0/35 | $<1.2 \times 10^{-6}$ |
| Wild-type | *Tc1::sfGFP* | 27 | ND | 27441 | 0/27 | $<1.4 \times 10^{-6}$ |
| *hrpc-1(gg910)* | *Tc1::sfGFP* | 36 | ND | 11704 | 0/36 | $<2.4 \times 10^{-6}$ |
| *rde-3(gg1160)* | *Tc1::sfGFP* | 41 | ND | 9150 | 4/41 | $1.1 \times 10^{-5}$ |
| *hrpc-1(gg910); rde-3(gg1161)* | *Tc1::sfGFP* | 42 | ND | 849 | 9/42 | $2.8 \times 10^{-4}$ |

\**gg1160* and *gg1161* are equivalent to *ne298*

Second, *rde-3* and *mut-16* limit *Tc1* activity in the germline of *C. elegans* [36,38]. *rde-3(-); Tc1::mScarlet-I* and *mut-16(pk710); Tc1::mScarlet-I* animals exhibited a >1000 fold increase in the rate of *Tc1* excision from *Tc1::mScarlet-I* in the germline as indicated by the appearance of animals expressing mScarlet-I in all cells (Table 2 and Figure 1C). Thus, excision of *Tc1* from *Tc1::mScarlet-I* is regulated by factors that regulate endogenous *Tc1* elements in the germline. Additionally, RDE-3 is reported to suppress excision of *Tc1* in the germline, but not in the soma [36]. We observed that RDE-3(-) animals exhibited increased *Tc1* excision from *Tc1::mScarlet-I* in the germline (see above) and did not show increase rates of *Tc1* excision from *Tc1::mScarlet-I* in the soma (Table 1).

Third, 78% of the time endogenous *Tc1* elements, which insert into TA dinucleotides, leave behind a four base pair TATGTA or TACATA excision footprint when excised from somatic cells [19]. We cloned and sequenced eleven independent somatic *Tc1::mScarlet-I* excision events observed in *Tc1::mScarlet-I* animals and in 8/11 cases (73%), we observed a TATGTA or TACATA four base pair insertion at the TA dinucleotide formerly occupied by *Tc1* in *Tc1::mScarlet-I* (Figure 1D).

Finally, we constructed a second *Tc1* reporter gene in which a *Tc1* element was inserted into the coding sequence of a superfolder green fluorescent protein (*sfgfp)* gene driven under the control of a distinct, ubiquitous promoter (*rpl-28*p) (Figure 1E). The reading frame of *Tc1::sfgfp* was engineered such that a typical somatic *Tc1* excision footprint (+4) would enable in-frame sfGFP translation (Figure 1E). *Tc1::sfgfp* behaved like *Tc1::mScarlet-I*, exhibiting properties previously ascribed to endogenous *Tc1* elements (Figure 1E). These properties included elevated germline *Tc1* excision rates in *rde-3(-)* animals (Table 2). For unknown reasons, the germline excision rate from *Tc1::sfgfp* in *rde-3(-)* animals occurred at a ≅10x lower rate than that of *Tc1::mScarlet-I* (Table 2). Together, these data establish that *Tc1* reporter genes recapitulate known properties of endogenous *C. elegans Tc1* elements and, therefore, can be used to explore transposon biology in an intact animal system.

### Identification of factors silencing *Tc1* in the germline

*C. elegans* RDE-3 and MUT-16 prevent *Tc1* mobilization in the germline [21,38]. *rde-3* encodes a poly(UG) nucleotidyltransferase [39,40] and *mut-16* encodes a low complexity Q/N-rich protein [38]. Both RDE-3 and MUT-16 localize within germ cells to non-membrane enclosed cytoplasmic organelles termed *Mutator* foci (Figure 2A), where they contribute to the biogenesis of endogenous (endo) small interfering (si)RNAs, which are thought to negatively regulate the *Tc1* transposase mRNA, thereby preventing *Tc1* transposition [41–44]. Using our *Tc1::mScarlet-I* reporter gene, we observed, as expected, that *mut-16* suppresses germline *Tc1* excision from *Tc1::mScarlet-I* in the germline (Table 2 and Figure 2B). Next, we used the *Tc1::mScarlet-I* reporter to ask if other siRNA-related genes and pathways might coordinate with RDE-3 *and* MUT-16 to silence transposons in the *C. elegans* germline. *Mutator* foci localize near the outer membrane of germ cell nuclei juxtaposed to nuclear pores and directly adjacent to two other perinuclear subcompartments of the germ granule termed the P granule and the Z granule [44,45] (Figure 2A). Animals lacking DEPS-1, which is a low-complexity protein that localizes to P granules and which contributes to P granule formation [46], exhibited an increase in germline *Tc1* excision from *Tc1::mScarlet-I* (Figure 2B, Table 2). Animals lacking the helicase ZNFX-1, which localizes to Z granules-where it binds mRNAs undergoing siRNA-based gene silencing [47] - exhibited a moderate increase in germline *Tc1* excision, which became evident after lineages were grown to high densities for two to three generations (Figure 2C, Table 2). The data suggests that the various perinuclear germ granule compartments help coordinate anti-transposon defenses in *C. elegans*, perhaps by concentrating siRNA pathway proteins near nuclear pores to surveille transposon mRNAs exiting nuclei [48]. In addition to these cytoplasmic functions, *C. elegans* siRNAs also regulate gene expression within nuclei by modifying chromatin states and inhibiting transcription elongation (termed nuclear RNAi) (Figure 2A) [49,50]. We asked if the nuclear RNAi pathway might contribute to transposon silencing in the germline. Indeed, animals lacking NRDE-2, which is an RNA binding protein required for nuclear RNAi [51] exhibited a moderate increase in germline *Tc1* excision, which was evident after two to three generations of growth (Figure 2C, Table 2). We conclude that the cytoplasmic and nuclear branches of the RNAi system coordinately limit transposon excision in the *C. elegans* germline.

**Figure 2.**
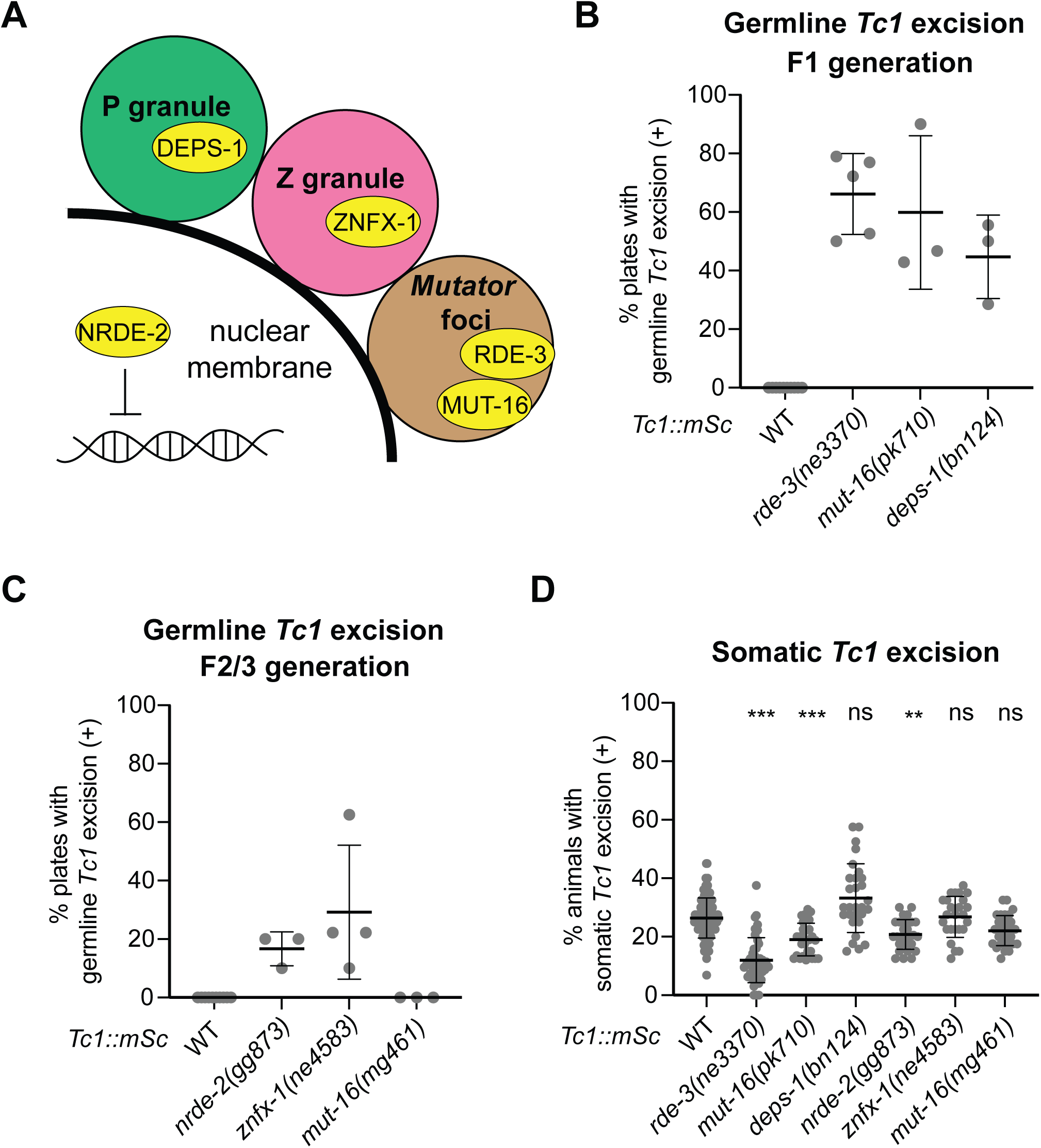
Germline and soma employ different mechanisms to regulate *Tc1* excision. **(A)** Schematic of the *C. elegans* RNAi pathway showing factors assayed for roles in suppressing *Tc1* excision, including the germ granule components DEPS-1 (located in the P granule), ZNFX-1 (in the Z granule), RDE-3 and MUT-16 (in *Mutator* foci), and the nuclear RNAi factor NRDE-2, which inhibits gene expression at the level of transcription. (**B** and **C**) Percentage of plates containing one or more animals with germline *Tc1* excision (indicated by mScarlet-I expression in all cells) when populations started with a single animal were expanded to the F1 generation **(B)** or F2/F3 generations **(C)**. Total plates scored per genotype per replicate were 6-19. ≥3 replicates scored per genotype. Error bars denote mean ± standard deviation (SD). WT, wild-type. **(D)** Somatic *Tc1* excision rates visualized as percentage of progeny animals expressing mScarlet-I in one or more somatic cells from independently singled animals. Data points represent independent lineages and are from ≥3 replicates. Total lineages scored per genotype were 25-116. WT is wild-type. ns, not significant; **, p≤ 0.01; ***, p ≤ 0.001 (Kruskal-Wallis test with Dunn’s test for multiple comparisons). Error bars denote mean ± SD.

### Germline anti-transposon systems do not regulate *Tc1* in the soma

RDE-3, MUT-16, and NRDE-2, which silence transposons in the germline (see above), are expressed and/or are active in both the germline and the soma [40,43,51]. Therefore, we asked if these inhibitors of germline *Tc1* might also inhibit *Tc1* excision in the soma. Surprisingly, presumed null mutations in *rde-3*, *deps-1*, *znfx-1*, or *nrde-2* did not increase *Tc1* excision in the soma (Figure 2D, Table 1) as they did in the germline. For unknown reasons, *rde-3*, *mut-16(pk710)*, and *nrde-2* mutants actually exhibited a subtle, yet statistically significant, decrease in somatic excision of *Tc1* from the *Tc1::mScarlet-I* reporter gene (Figure 2D). The *mut-16(pk710)* allele ablates MUT-16 function in both the soma and germline while the *mut-16(mg461)* allele deletes a portion of the *mut-16* promoter, which abolishes MUT-16 function in the soma, but not the germline [43]. While *mut-16(pk710)* animals showed elevated rates of *Tc1* excision in the germline, somatic excision rates were not elevated (Figure 2D, Table 1). The soma-specific allele *mut-16(mg461)* failed to alter excision rates in either the germline or the soma (Figures 2C-D, Tables 1-2). Together, the data show that the germline RNAi-based anti-transposon system does not limit *Tc1* excision in the *C. elegans* soma, despite components of these pathways being present and active in this tissue.

### Identification of genes that limit *Tc1* excision in the soma

We wondered if other, currently unknown systems might regulate *Tc1* in the soma. To identify such systems, we conducted a forward genetic screen using *Tc1::mScarlet-I; Tc1::sfgfp* double reporter animals to identify mutations that increased the frequency of *Tc1* excision from both reporters in somatic cells (Figure S1A). The screen identified 21 mutant strains exhibiting >3x more somatic cells expressing mScarlet-I and sfGFP than wild-type animals. Seven of these 21 mutant strains exhibited a dramatic >60x elevation in somatic *Tc1* excision as well as a delay in developmental timing (see below). We focused our efforts on characterizing these seven alleles due to the high expressivity of their elevated *Tc1* excision phenotypes and because their shared developmental pleiotropy hinted at shared molecular function.

### HRPC-1 and STM-1 limit *Tc1* excision in the soma

Positional mapping and whole genome sequencing identified candidate mutations in all seven strains. In six of the seven strains, we identified mutations in the gene <u>z</u>inc finger putative transcription factor (*ztf)-4* (Figure 3A), which were each associated with a ≅85x elevation in *Tc1* somatic excision (mean number of mScarlet-I positive cells per animal: 0.44 in wild-type and 37.4 in mutant) (Figure 3B). *C. elegans ztf-4* is predicted to encode one zinc finger and an RNA recognition motif (RRM) domain (Figure 3A). Basic Local Alignment Search Tool (BLAST) analyses suggested that ZTF-4 is orthologous to vertebrate heterogeneous nuclear ribonucleoprotein C (HNRNPC) protein, with homology most pronounced in the RRM domain (Figure S1B). For clarity, we renamed *ztf-4* as <u>h</u>ete<u>r</u>ogeneous nuclear <u>p</u>rotein <u>C</u>-like 1 (*hrpc-1)*. All six *hrpc-1* mutations identified by the genetic screen encoded missense alleles, four of which altered amino acids within the RRM domain of HRPC-1 (Figures 3A and S1B). CRISPR/Cas9-based mutagenesis introducing the A132V RRM mutation into wild-type *hrpc-1* resulted in animals that exhibited elevated levels of somatic *Tc1* excision from *Tc1::mScarlet-I* and *Tc1::sfgfp*, confirming that *hrpc-1* mutation causes an enhanced somatic *Tc1* excision phenotype (Figure 3C). CRISPR/Cas9-based deletion of >90% of the *hrpc-1* locus also resulted in animals exhibiting high levels of somatic *Tc1* excision (Figure S2), indicating that one function of HRPC-1 in *C. elegans* is to limit *Tc1* excision in somatic cells, and suggesting that the six *hrpc-1* alleles identified in our genetic screen were loss-of-function.

**Figure 3.**
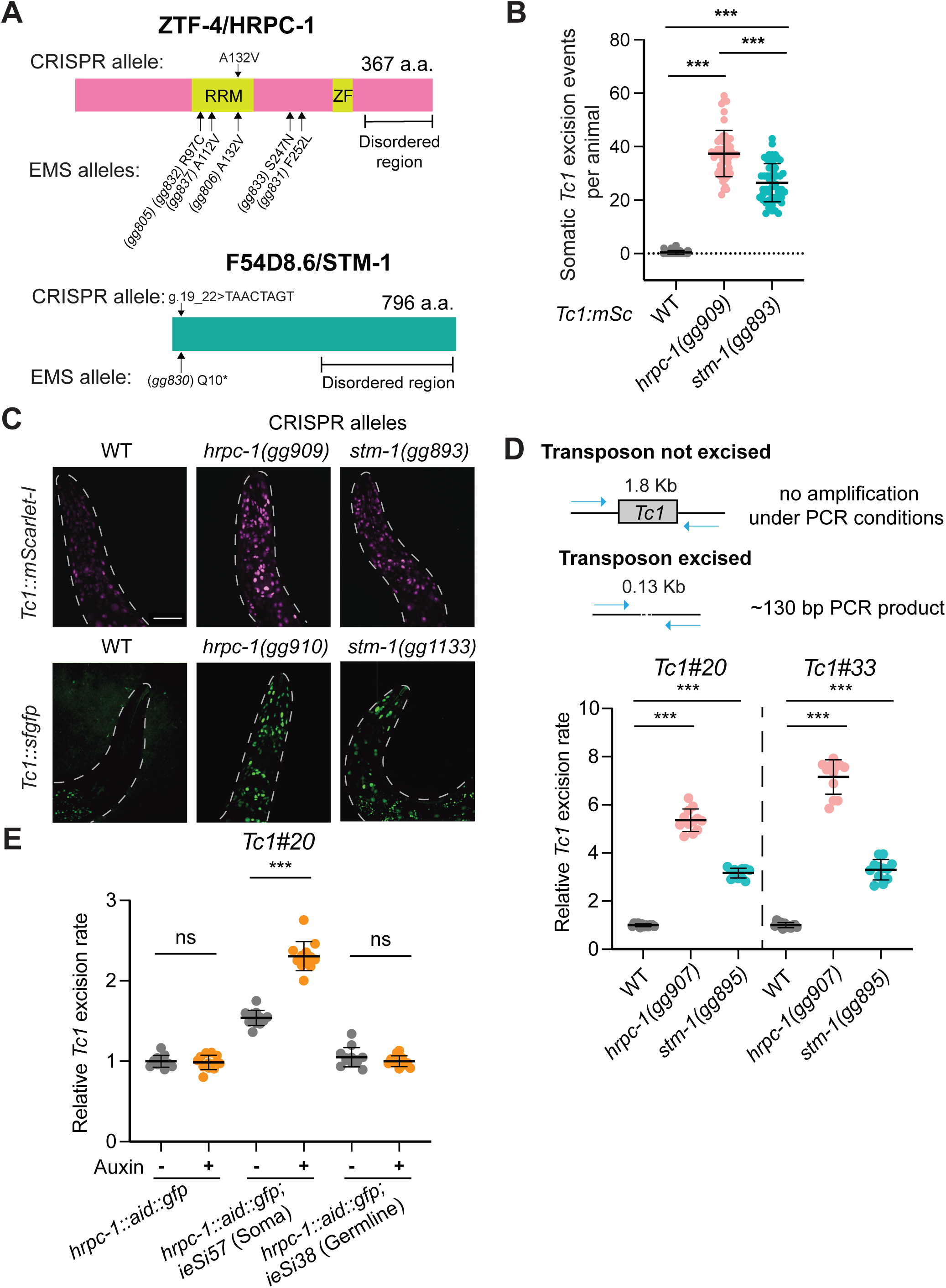
HRPC-1 and STM-1 regulate *Tc1* primarily in the soma. **(A)** *ztf-4*/*hrpc-1* and *F54D8.6*/*stm-1* alleles identified in the genetic screen (“EMS allele(s)”) or generated by CRISPR-Cas9 (“CRISPR allele”) are shown. Unless otherwise indicated, *hrpc-1* and *stm-1* alleles used in this work are the CRISPR-Cas9-derived mutations that lead to HRPC-1(A132V) or STM-1(g.19_22>TAACTAGT). a.a., amino acid; RRM, RNA recognition motif; ZF, zinc finger domain. **(B)** Quantification of somatic excision events ((mScarlet-I (+) somatic cells)) in *Tc1::mScarlet-I* expressing strains of the indicated genotypes. n=51-59 animals per genotype. Data are from 3 biological replicates. ***, p ≤ 0.001 (Kruskal-Wallis test with Dunn’s test for multiple comparisons). Error bars denote mean ± SD. WT, wild-type. **(C)** Fluorescent micrographs of the anterior (head) of *Tc1::mScarlet-I* or *Tc1::sfgfp* animals of indicated genotypes showing that the number of sfGFP or mScarlet (+) cells is elevated in strains of the indicated genotypes. Scale bar, 30 μm. **(D)** (Top) Schematic of qPCR-based assay to quantify endogenous *Tc1* excision events from genomic DNA. qPCR settings were optimized such that amplicons are detected only if *Tc1* is excised. (Bottom) Quantification of *Tc1* excision in *hrpc-1* or *stm-1* for two endogenous *Tc1* sites (*Tc1#20*: WBTransposon00000020; *Tc1#33*: WBTransposon00000033). Fold changes in excision rate were normalized to mean value in wild-type. n=12 per genotype, from 3 biological replicates. ***, p ≤ 0.001 (one-way ANOVA with Dunnett’s multiple comparisons test). Mean ± SD. WT, wild-type. **(E)** qPCR quantification of *Tc1* excision for *Tc1#20* in strains with or without auxin-induced depletion of HRPC-1 in the germline or soma (see Methods). n=11-12 per genotype/treatment condition, from 3 biological replicates. ns, not significant; ***, *p* ≤ 0.001 (Student’s *t* test). Mean ± SD.

In the remaining mutant strain, which did not harbor a *hrpc-1* mutation, and which exhibited an ≅60x elevation in *Tc1* somatic excision (mean number of mScarlet-I positive cells per animal: wild-type, 0.44; mutant, 26.5) (Figure 3B), genome sequencing and positional mapping identified a candidate mutation in the gene *F54D8.6* (Figure 3A). CRISPR/Cas9-based introduction of a stop codon, or a short insertion that creates a frameshift at the 7th amino acid of *F54D8.6*, resulted in animals exhibiting high rates of *Tc1* excision from *Tc1::mScarlet-I* and *Tc1::sfgfp* in somatic tissues (Figure 3C). Based upon these results and results presented below, *F54D8.6* was named <u>s</u>uppressor of transposon <u>m</u>obilization (*stm*)*-1*. Because *stm-1(gg830)* encodes a premature (Q10) stop in the predicted *stm-1* open reading frame (Figure 3A), *stm-1(gg830)* is likely loss-of-function, indicating that a wild-type function of STM-1 is to limit *Tc1* excision. *stm-1* is predicted to encode a protein of unknown function, which lacks obvious homologs outside of *Caenorhabditae*.

We asked if, as expected, HRPC-1 and STM-1 regulate endogenous *Tc1* elements. We used qPCR to quantify rates of *Tc1* excision from two of the ≅30 endogenous *Tc1* elements present in the *C. elegans* N2 Bristol strain genome (Figure 3D). The analysis revealed a 3-8 fold increase in *Tc1* excision events at these endogenous *Tc1* loci in both *hrpc-1* or *stm-1* mutant animals, confirming that HRPC-1 and STM-1 are regulators of endogenous *Tc1* elements (Figure 3D).

### The *C. elegans* soma and germline utilize different transposon regulatory strategies

We used our reporter strains to assess the rate of germline *Tc1* excision in *hrpc-1* and *stm-1* animals. Interestingly, *hrpc-1* and *stm-1* animals did not exhibit detectable increases in germline *Tc1* excision from *Tc1* reporter genes, as was apparent in somatic cells in these animals (Table 2). The data suggests that loss of HRPC-1 or STM-1 does not increase *Tc1* excision rates in the germline at least to levels detectable in these assays (see below). To test the idea that HRPC-1 and STM-1 act primarily in the soma to regulate *Tc1*, we used an auxin-degron system to deplete a degron::GFP tagged HRPC-1 from either the germline or the soma. Fluorescent imaging confirmed auxin-dependent depletion of HRPC-1::AID::GFP from the soma or the germline (Figure S3). Depletion of HRPC-1 from somatic tissues resulted in an increase in *Tc1* excision (Figure 3E). Note that in these experiments, animals that were not treated with auxin expressed less HRPC-1::AID::GFP (Figure S3), and showed *Tc1* excision rates greater than control animals not expressing somatic TIR1 (Figure 3E), suggesting that some HRPC-1::AID::GFP depletion occurs even in the absence of auxin treatment. Depletion of HRPC-1::AID::GFP from the germline, however, did not impact *Tc1* excision (Figure 3E). Similarly, ablation of germ cells, using a temperature sensitive *glp-1(q224)* allele in either *hrpc-1 or stm-1* animals did not decrease endogenous *Tc1* excision rates (Figure S4). These data show that loss of HRPC-1 results in an increase in *Tc1* excision in the soma, but not in the germline, at least to levels detectable in these assays (see below).

To assess if HRPC-1 might regulate *Tc1* excision in the germline at levels below the detection threshold of our assays, we asked if loss of HRPC-1 would enhance germline *Tc1* excision in animals lacking the germline RNAi factor RDE-3. To address this question, we assayed germline *Tc1* excision frequencies in *rde-3* and *rde-3; hrpc-1* double mutant animals. The analysis revealed a >10 fold increase in the germline *Tc1* excision rates in *rde-3; hrpc-1* mutants compared to *rde-3* single mutants for *Tc1::sfgfp* (Table 2) and a 2x increase in excision of an endogenous *Tc1* element (Figure S5). The data suggests that HRPC-1 acts in both the soma and germline to prevent *Tc1* excision, however, the consequences of losing HRPC-1 in germ cells are minor in animals harboring an intact RNAi system.

### HRPC-1 and STM-1 are ubiquitously expressed nuclear proteins that interact *in vivo*

To identify where and when *hrpc-1* and *stm-1* are expressed, we introduced GFP or mScarlet-I tags to the *hrpc-1* and *stm-1* genes, respectively. Wild-type animals develop from egg to larval stage four (L4) in ≅48 hours. *hrpc-1* and *stm-1* mutant animals exhibit delays in larval development; these animals took ≅72 and 60 hours respectively to reach the L4 stage of development (Figure S6). Epitope-tagged *hrpc-1::gfp* or *stm-1::mScarlet-I* expressing animals did not exhibit delays in larval development, indicating that the fluorescent tags introduced into these genes did not impact protein function, at least with regards to developmental timing (Figure S6). Fluorescent microscopy revealed that HRPC-1::GFP and STM-1::mScarlet-I were expressed in most or all cells of the soma (Figure 4A) and the germline (Figure 4B), and in these cells, HRPC-1::GFP and STM-1::mScarlet-I localized predominantly within nuclei (Figures 4A-B). We conclude that HRPC-1 and STM-1 are ubiquitously expressed nuclear proteins. The shared expression patterns of HRPC-1 and STM-1, and the shared developmental phenotypes of *hrpc-1* and *stm-1* mutants, hints that HRPC-1 and STM-1 might interact physically. To test this idea, we introduced hemagglutinin (HA) epitopes to the *hrpc-1* gene for immunoprecipitation mass spectrometry (IP-MS) experiments. *hrpc-1::3xha* animals did not exhibit delays in larval development, suggesting the HA epitopes do not negatively impact HRPC-1 function (Figure S6). IP-MS of HRPC-1::3xHA identified STM-1 as the most enriched protein in HRPC-1::3xHA precipitates (Fig. 4C-D, Figure S7, and Supplementary Table 1), suggesting that HRPC-1 and STM-1 associate physically *in vivo*.

**Figure 4.**
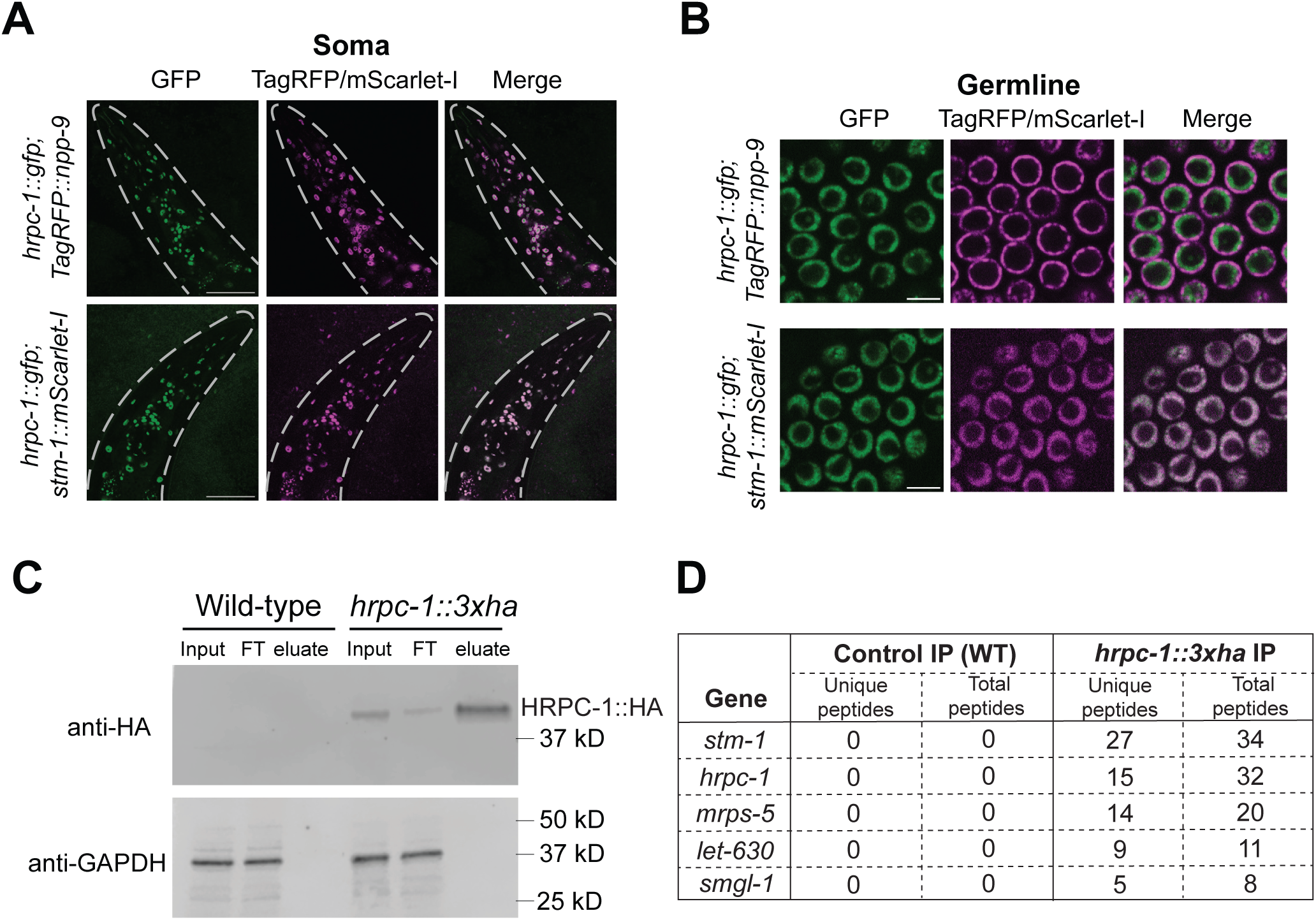
HRPC-1 and STM-1 localize to the nucleus and interact *in vivo*. **(A and B)** Representative confocal images showing expression of HRPC-1::GFP in the soma (A) and germline (B). (Top rows) Nuclei are marked with the nuclear pore protein TagRFP::NPP-9. (Bottom rows) STM-1::mScarlet-I colocalizes with HRPC-1::GFP in the nucleus. Scale bar, 40 μm (**A**), 5 μm (**B**). **(C** and **D)** IP-MS experiment to identify HRPC-1 interactors. (**C**) Gel image showing immunoblotting of HA and GAPDH in indicated samples from the IP-MS experiment. Wild-type strain is the control. FT, flow-through fraction. (**D**) Number of unique and total peptides of the top 5 proteins that were only identified from *hrpc-1::3xha* IP-MS. Results of another independent replicate are shown in Figure S7. In the independent IP-MS replicate, STM-1 was the most common co-precipitating protein and the other 3 proteins did not reproduce. WT, wild-type.

### HRPC-1/STM-1 associate with *Tc1*-containing RNAs

Because HRPC-1 possesses an RRM RNA binding domain, we wondered if HRPC-1 might interact with *Tc1* RNA. To test this idea, we used *hrpc-1::3xha* animals to conduct anti-HA immunoprecipitation (IP) and then RT-qPCR to quantify co-precipitating RNAs. The analysis showed that IP of HRPC-1::3xHA enriched *Tc1* RNA in the precipitate >300x over levels of *Tc1* RNA observed in control IPs, which employed animals not expressing HRPC-1::3xHA (Figures 5A-B). HRPC-1::3xHA IP did not enrich a randomly selected mRNA (*cdc-42*), which does not contain a *Tc1* element (Figure 5B), suggesting that the interaction of HRPC-1 with the *Tc1* RNA is at least somewhat specific. Introduction of the A132V RRM variant into HRPC-1::3xHA by CRISPR negatively impacted the expression of HRPC-1::3xHA (see Figure 5A-input, S8A-input, Figure S8C). Nonetheless, when levels of *Tc1* RNA in the pellet were normalized to levels of immunoprecipitated HRPC-1 protein, HRPC-1(A132V)::3xHA failed to enrich *Tc1* RNA, suggesting that the A132C RRM mutation disrupts HRPC-1/*Tc1* RNA interactions (Fig. 5B and S8B). We next asked if STM-1 might be required for HRPC-1::3xHA to associate with *Tc1*-containing RNAs. Again, HRPC-1 was poorly expressed in *stm-1* mutant backgrounds (see Figure 5A-input, S8A-input). Nonetheless, when levels of RNA in the pellet were normalized to levels of immunoprecipitated HRPC-1 protein, HRPC-1::3xHA co-precipitated with less *Tc1* RNA in an *stm-1* mutant than in *stm-1(+)* animals, although HRPC-1::3xHA retained some ability to associate with *Tc1* in the absence of STM-1 (Fig. 5B and S8B). The data suggest that HRPC-1 associates with *Tc1*-encoded RNA and that this association depends, at least in part, on STM-1 and the RRM domain of HRPC-1. The data also support the idea that HRPC-1 and STM-1 act together in a pathway to repress *Tc1*.

**Figure 5.**
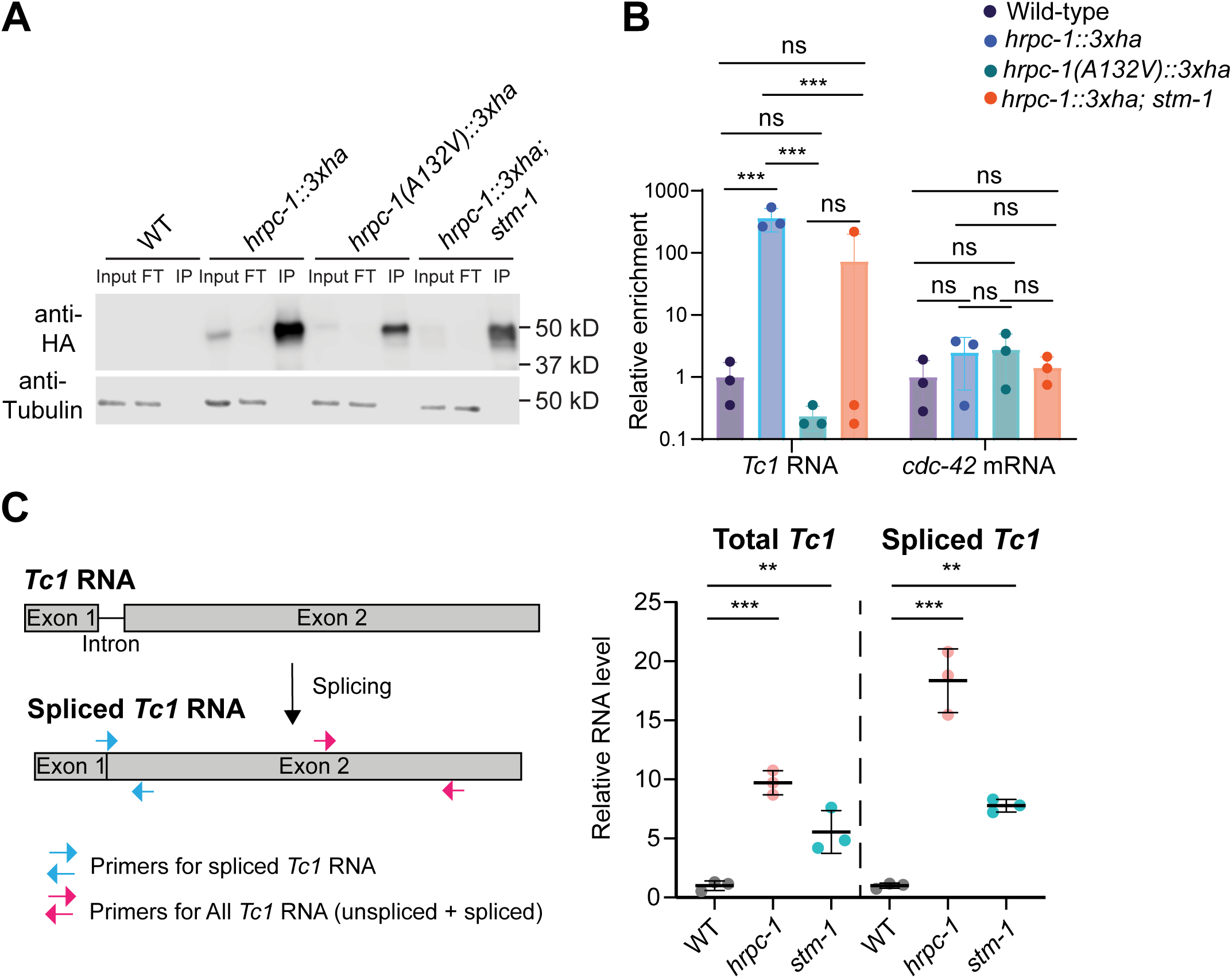
HRPC-1 binds *Tc1* RNA and suppresses *Tc1* RNA levels. **(A and B)** RNA immunoprecipitation experiment for detection of HRPC-1 interaction with *Tc1* RNA. **(A)** Representative image of HA immunoblotting of indicated proteins (tubulin, loading control). FT, flow-through fraction. IP, immunoprecipitation fraction. WT, wild-type. **(B)** RT-qPCR quantification of indicated RNAs in HRPC-1::HA immunoprecipitates. Levels of RNA in immunoprecipitates were normalized to levels of RNA in inputs. Mean enrichment level of wild-type sample was set as 1. ns: not significant; ***, *p* ≤ 0.001 (Two-way ANOVA with Šídák’s multiple comparisons test.) Data are from 3 biological replicates. Mean ± SD. **(C)** (Left) Primers for RT-qPCR of total *Tc1* (includes unspliced and spliced *Tc1*) or spliced *Tc1* RNA as indicated. The forward primer for detecting spliced *Tc1* RNA spans the *Tc1* exon-exon junction. (Right) RT-qPCR quantification of total *Tc1* and spliced Tc1 RNA levels in *hrpc-1* and *stm-1*. **, p ≤ 0.01; ***, p ≤ 0.001 (one-way ANOVA with Dunnett’s multiple comparisons test.). Data are from 3 biological replicates. Mean ± SD. WT, wild-type.

### HRPC-1/STM-1 regulate *Tc1*-containing RNAs

Since HRPC-1 associates with *Tc1* RNA, we wondered whether HRPC-1/STM-1 might regulate *Tc1* RNA abundance and/or processing. Indeed, we observed elevated (5-10x) levels of *Tc1* RNA in *hrpc-1* and *stm-1* mutant animals (Figure 5C, total *Tc1*), indicating that HRPC-1/STM-1 negatively regulates *Tc1* RNA. While conducting the above RT-PCR analysis, we noticed that the size of *Tc1* RNA differed in *hrpc-1* and *stm-1* mutants: In wild type animals, RT-PCR of *Tc1* generated a single PCR product while in *hrpc-1* and *stm-1* mutants the RT-PCR produced two bands, which differed in length by ∼40 base pairs (Figure 6). The *Tc1* RNA contains a single 41 nucleotide intron that must be spliced for the *Tc1* mRNA to be translated [52]. For unknown reasons (see discussion), most *Tc1* RNA is not spliced in wild-type *C. elegans* [53]. Sequencing revealed that the two PCR products observed in *hrpc-1* and *stm-1* mutants were spliced and unspliced *Tc1* RNAs (Figure S9). RT-qPCR analysis confirmed that *Tc1* RNA is more efficiently spliced in the absence of HRPC-1 and STM-1 (Figure 5C, spliced *Tc1*). These results suggest that HRPC-1 and STM-1 could suppress *Tc1* RNA levels- and *Tc1* activity-by inhibiting intron processing of *Tc1*, thereby decreasing *Tc1* RNA stability.

**Figure 6.**
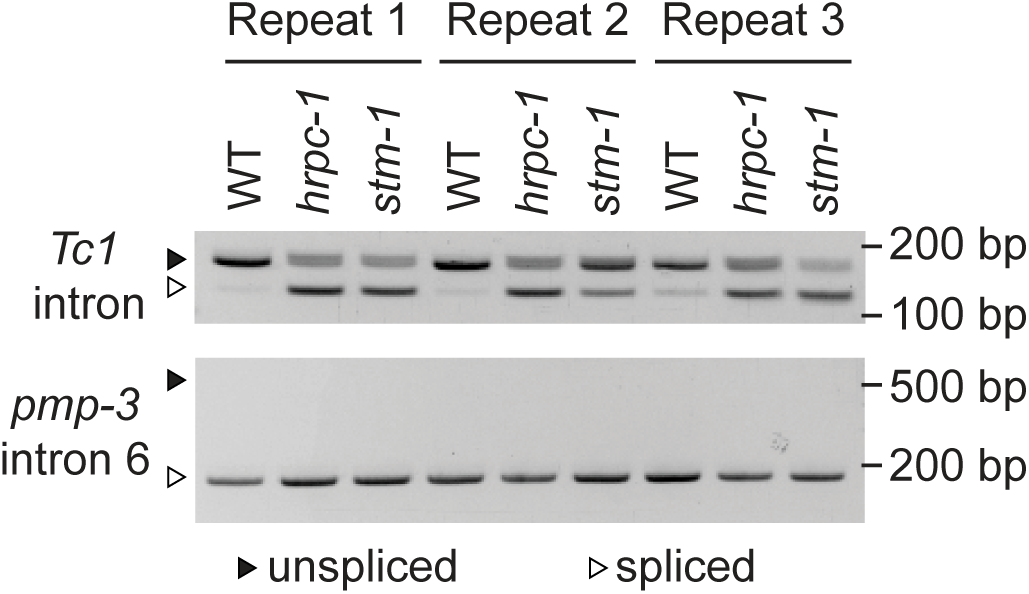
HRPC-1 and STM-1 affect splicing of *Tc1* RNAs. Electrophoresis of RT-PCR products showing *Tc1* intron splicing efficiency in animals of the indicated genotypes. Triangles indicate bands for unspliced (filled) and spliced (hollow) RNAs. Splicing of the *Tc1* intron is increased in *hrpc-1* and *stm-1* mutants compared to WT. Splicing of a control *pmp-3* intron is similarly efficient across all three genotypes. WT, wild type.

In summary, we report the development of fluorescence-based reporter genes for studying transposon regulation in a living animal. Using these tools, we show that *C. elegans* uses different systems to regulate *Tc1* excision in its germline and soma and we identify a pathway involving HRPC-1 and STM-1 that limits *Tc1* excision primarily in the soma, likely through binding *Tc1* RNA to limit *Tc1* intron splicing.

## Discussion

Here, we develop fluorescence-based reporter genes to visualize transposon excision at single-cell level resolution in a living animal. We use these reporter genes to show that *C. elegans* employs different strategies to regulate *Tc1* in its soma and germline. Consistent with previous research [54], we find that cytoplasmic factors that localize to- and/or help organize-germ granules (DEPS-1 and ZNFX-1) [46,47], and a nuclear factor that acts in the nuclear branch of the RNAi pathway (NRDE-2) [51], contribute to transposon suppression in the germline. The results reinforce the idea that the germline anti-transposon system is complex and multi-faceted, involving both cytoplasmic and nuclear branches.

Our results also show that, surprisingly, the germline RNAi pathway is not employed in the soma to regulate *Tc1*, despite many of the germline RNAi components being expressed in the soma, where they contribute to RNAi-based gene silencing [40,43,55]. Why wouldn’t *C. elegans* use the germline RNAi system to control transposons, such as *Tc1*, in somatic tissues? The RNAi pathway is complex, employing dozens of proteins subcompartmentalized into both membrane and non-membrane delimited organelles in germ cells [56]. It is possible that *C. elegans*, and perhaps other organisms, choose not to deploy RNAi to combat transposons in their somatic tissues because the fitness costs of somatic transposition are not as high in the soma as they are in the germline and because the system is too energetically costly to merit deployment. Rather, we find that *C. elegans* uses an HNRNPC/STM-1-based system to regulate *Tc1* in its soma. Interestingly, human HNRNPC binds and prevents exonization of *Alu* retrotransposons, which are the most abundant transposons in the human genome [26,27]. Thus, the role of HRPC-1 in regulating transposon-encoded RNAs may be conserved. Future studies are needed to assess if the role of HNRNPC in regulating transposons in nematodes and mammals reflects an ancient function of the HNRNPC proteins, or if the shared functions are due to convergent evolution. It will also be of interest to determine the breadth of transposon families regulated by HNRNPC-like proteins in eukaryotes.

Human HNRNPC is thought to bind antisense-oriented *Alu* RNAs, at least in part, by recognizing single-stranded poly-U tracts present in *Alu* RNAs synthesized from genes containing antisense, intronic *Alu* insertions [27]. By recognizing poly-U tracts, HNRNPC prevents binding of the splicing factor U2AF65, thereby preventing *Alu* elements from becoming exonized into host mRNAs [27]. It is possible that *C. elegans* HRPC-1 may detect the *Tc1* RNA by binding poly-U tracts-the *Tc1* RNA contains 19 poly-U tracts ≥5 base pairs. Additionally, RNA secondary structure can influence HNRNPC interaction with RNA [57,58] and transposon-encoded RNAs often adopt secondary structures [26,41], which may also serve as a recognition feature for HRPC-1/HNRNPC. Understanding how HRPC-1 and HNRNPC identify transposon-encoded RNAs for neutralization will be an important question for future studies to address.

We find that splicing of the *Tc1* intron is more efficient in *hrpc-1* and *stm-1* mutant animals, which would be expected to increase transposase expression and, therefore, *Tc1* mobilization. Intriguingly, splicing of the third intron of the DNA transposon *P* element in *Drosophila* is also actively regulated, and this splicing determines whether the translated protein is active as a transposase [59]. In *Drosophila* somatic cells, splicing of *P* elements is repressed by RNA-binding proteins that recognize sequence specific elements upstream of the alternatively spliced third intron [60]. Taken together with our observations, the data hint that the regulation of transposon splicing could be a common strategy employed by animal cells to regulate DNA transposons in the soma. A final question emerging from our work is why would *C. elegans Tc1* and human *Alu* possess signals that allow HRPC-1/HNRNPC to bind and regulate their splicing. Transposons cannot survive without their hosts, and transposition in the soma does not benefit the transposon. Therefore, we speculate that transposons are under selective pressure to evolve sequences that allow HRPC-1/HNRNPC (and perhaps other systems) to limit their splicing in the soma in order to limit transposon exonization into host genes, thereby minimizing the fitness costs incurred by both host and parasite. Asking if *C. elegans* HRPC-1 prevents the exonization of *Tc1*, or other transposons, into host mRNAs will be a strong test of this latter idea.

## Materials and Methods

### Strains

All strains were cultivated on standard Nematode Growth Medium (NGM) plates and maintained at 20°C with *Escherichia coli* strain OP50 as food source unless otherwise noted [62]. *glp-1*(*q224ts*) animals were maintained at 15°C. CRISPR strains were made with the CRISPR/Cas9 strategies described by Arribere et al., or Ghanta and Mello [63,64]. Guide RNAs were chosen according to the CRISPOR.org tool [65]. *Tc1::mScarlet-I* and *Tc1::sfgfp* reporter strains were made via CRISPR/Cas9 with Saptrap [66] combined with self-excising cassette selection strategy [66,67]. List of strains and details is available in Supplementary Table 2.

### *Tc1* reporter plasmids construction

*Tc1::mScarlet-I* and *Tc1::sfgfp* reporter plasmids for CRISPR/Cas9 injection to make *Tc1* reporter strains were constructed by using Saptrap to assemble components into the pDD379 vector [66]. *Tc1::mScarlet-I* reporter plasmid donor sequence consists of *Tc1* (1613 base pairs, including a TA dinucleotide at the 3’ end) inserted after the TA dinucleotide at position -7 to -8 of the 622 base pair *eft-3* promoter, followed by *mScarlet-I* sequence with nuclear-localization sequences (NLS) (SV40 NLS at 5’ end; *egl-13* NLS at 3’ end) and the *tbb-2* 3’UTR. *Tc1::sfgfp* reporter plasmid donor sequence consists of a 523 base pair *rpl-28* promoter, followed by a start codon and an 18 base pair flexible linker before the Tc1 sequence (1615 base pairs, including TA dinucleotide at both 5’ and 3’ ends).

The *Tc1* sequence is followed by another 18 base pair flexible linker before the *sfgfp* sequence with NLSs (SV40 NLS at 5’ end; *egl-13* NLS at 3’ end of *sfgfp*) and the *unc-54* 3’UTR. The nuclear localized *sfgfp* sequence lacks a start codon. The *Tc1* sequence in *Tc1::mScarlet-I* and *Tc1::sfgfp* plasmid constructs is the sequence of *Tc1* with WormBase ID WBTransposon00000015, which does not contain any polymorphisms [11]. *Tc1::mScarlet-I* reporter plasmid guide RNA and homology arm components were designed to insert the reporter at Chr I: -5.32 cM. *Tc1::sfgfp* reporter plasmid guide RNA and homology arm components were designed to insert the reporter at Chr III: -0.85 cM. These sites were chosen according to insertion sites established for Mos1 mediated single copy insertion (MosSCI) that allow germline expression of inserts [68,69].

### Microscopy

In Figures 1B, 1C, 1E images, animals were immobilized with 0.05% sodium azide and mounted on glass slides; in Fig S3 images, animals were mounted on glass slides without immobilization. Images in Figures 1B, 1C, 1E, S3 were taken using the Axio Observer.Z1 fluorescent microscope (Zeiss) with the Plan-Apochromat 20x/0.8 M27 objective or the Plan-Apochromat 63x/1.4 Oil DIC M27 objective. Images were acquired using ORCA-Flash4.0 V2 digital camera (Hamamatsu) and ZEN 2 (blue edition) software (Zeiss). For Figures 3C, S2, and 4A-B images, larval stage 4 (L4) animals (Figures 3C and S2) or one day old adult animals (Figures 4A-B) were immobilized with polystyrene beads (Polysciences, Cat #00876) and mounted on 10% agarose pads. Images were taken using the Dragonfly 505 spinning-disk confocal microscope with iXon Ultra 888 EMCCD camera (Andor Technologies) using Nikon 60x/1.40 (Figure 3C, S2, 4A) or 100x/1.45 (Figure 4B) oil immersion objectives. Images were processed using Fiji [70]. Microscope images in Figures 1B, 1C, 3C and S2 were stitched together from two or more images because a single image was not able to capture the full x-y dimension. Figure 3C and S2 images are maximum intensity projections of z-stack images taken at a step size of 0.5 μm spanning the thickness of the animal.

### *Tc1* excision footprint analysis

*Tc1::mScarlet-I*; *glp-1*(*q224ts*) strains were bleach-synchronized and grown from embryonic stage at 25°C for 3 days to reach adult stage. Adult worms were washed off the plate with M9 buffer and lysed with worm lysis buffer (50mM KCl, 10mM pH8.3 Tris, 2.5mM MgCl^2^) with proteinase K (0.2 mg/ml). Lysates were used as templates for PCR reactions using primers: F: 5’-CTACCGTCCGCACTCTTC-3’; R: 5’-CTTACGCTTCTTCTTTGGC-3’. PCR products were run on an agarose gel, and band size around 100 base pairs was excised and gel purified (Qiagen #27206). Purified PCR products were TA-cloned using the pGEM-T Easy Vector System (Promega, A1360), according to the manual instructions. White transformants were isolated, grown in liquid culture, and miniprepped. Plasmids were sent for Sanger sequencing using the primer M13F-40: 5’-GTT TTC CCA GTC ACG AC-3’.

### Soma *Tc1* excision rate quantification

For the soma *Tc1* excision assay in Table 1, for all genotypes except *hrpc-1* and *stm-1*, L4 animals (P0) were singled onto 6 cm NGM plates and kept at 20°C for their progeny animals to grow. 4-5 days later, each plate was examined under the AxioZoom.v16 fluorescence microscope (Zeiss) with the ApoZ 1.5x/0.37 objective. Progeny animals (F1) of the original singled animal were visually scored for presence or absence of somatic cells with mScarlet-I expression, to determine the number of progeny animals on a single plate that had somatic mScarlet-I expression. All visual scoring of soma mScarlet-I expression was done with the examiner blinded to genotype. This data is also shown as Figure 2D. For *hrpc-1* and *stm-1*, percentage of animals showing somatic cell mScarlet-I (+) was 100%, and L4 stage animals were examined under the Dragonfly 505 spinning-disk confocal microscope using the Nikon 60x/1.40 oil immersion objective to quantify number of somatic cells with mScarlet-I expression per animal. This data is also shown as Figure 3B.

### Germline *Tc1* excision rate quantification

Previous labs have established a Poisson distribution method for estimating germline transposon excision rate in *C. elegans* [19,36,37,71]. This method relies on establishing single lineages of animals, which are then scored as a population for whether or not excision event(s) have occurred at all in the span of the propagation time. Taking into account the amount of animals present in each lineage at time of scoring and the proportion of lineages that have excision event(s), a germline excision rate is calculated using the Poisson distribution formula [19,36]: f = - (ln(N/T))/n, f: germline Tc1 excision frequency, T: total number of lineages scored, N: total number of lineages without germline *Tc1* excision events, n: average number of animals per lineage. This calculation method addresses the concern of counting a single germline *Tc1* excision event multiple times, since a single animal that undergoes a germline excision event will generate many progeny presenting with germline excision events that result from inheriting the excised *Tc1* locus instead of undergoing actual independent germline excision events (jackpot effect) [19,36,37,71].

In our germline excision experiments (Table 2), L4 animals were singled onto 6 cm NGM plates and allowed to propagate at 20°C. 20x concentrated OP50 was added to plates during the interval of growth. After several days of propagation, when plates have reached either F1 or F2/3 generations, each plate (i.e. each lineage) was scored for presence or absence of germline *Tc1* excision event(s). Scoring was done under the AxioZoom.v16 fluorescence microscope (Zeiss) with the ApoZ 1.5x/0.37 objective for *Tc1::mScarlet-I* background animals and under the Leica M165 FC with a Plan APO 1.0x objective for *Tc1::sfgfp* background animals. A germline excision event is indicated by presence of an animal in which all cells are expressing mScarlet-I or sfGFP. The generation at which animals are scored for *Tc1* excision (F1 or F2/3) varies with genotype and depends on how frequent germline *Tc1* excision is in that particular genotype. The number of animals on each plate at time of scoring was estimated by counting animals in one region of the plate, then multiplying by plate area. The average number of animals on each plate (“n” variable in the Poisson distribution formula) is calculated by summing the number of animals on each plate for a genotype, then dividing this sum by the total number of plates for that genotype. Germline *Tc1* excision rates were calculated using the Poisson distribution formula. In genotypes where no germline *Tc1* excision event was detected, germline *Tc1* excision rates were indicated as less than the rate that would have been if one plate of all plates scored had germline *Tc1* excision event(s). All visual scoring was done with the examiner blinded to genotype.

### Forward genetic screen for *Tc1* excision regulators

*Tc1::mScarlet-I; Tc1::sfgfp* strains were treated with 47 mM ethyl methanesulfonate for four hours at 23°C (P0 generation), then washed with M9 buffer. At the F2, F3, F4 generations, mutants were visually screened under the AxioZoom.v16 fluorescence microscope (Zeiss) and mutants that had a high number of somatic cells expressing mScarlet-I were singled onto new plates. After the singled mutants gave rise to progeny, their progeny animals were verified to have high levels of somatic mScarlet-I and somatic sfGFP using the Axio Observer.Z1 fluorescent microscope (Zeiss). The screen was done in two rounds; for the first round, ∼69,000 haploid genomes were screened, and mutants were picked at F2 and F3 generations; for the second round, ∼180,000 haploid genomes were screened, and mutants were picked at F2, F3 and F4 generations. Mutant genomic DNA was extracted and sent for whole genome sequencing (Biopolymers Facility, Harvard Medical School). Based on whole genome sequencing and positional mapping information, causal alleles were identified.

### *Tc1* excision qPCR assay

L4 stage animals were lysed in worm lysis buffer (50mM KCl, 10mM pH8.3 Tris, 2.5mM MgCl_2_) with proteinase K (0.2 mg/ml). DNA lysates were used for qPCR assays using the iTaq Universal SYBR Green Supermix (Bio-Rad, 1725121), as instructed by the manual, and run on CFX Connect Real-Time PCR Detection System (Bio-Rad, 1855201). qPCR conditions were as follows: Initial denaturation at 95°C for 5 min followed by 40 cycles of denaturation at 95°C for 15 sec and annealing/extension/plate read at 61°C for 30 sec. Tc1 excision fold changes were calculated using the comparative Cq method [72]. Mean Cq value of wild-type strain was used as the baseline for calculating Tc1 excision fold change for each experiment. *eft-3* was used as the housekeeping gene. Primers used for qPCR analysis to detect *Tc1#20* and *Tc1#33* excision are listed in Supplementary Table 1.

### Multiple sequence alignment

Multiple sequence alignment was performed using MUSCLE v3.8.31 in Jalview v2.11.4.1 [73–75]. Protein sequences used for alignment were imported from Uniprot [76].

### Developmental timing assay

Ten gravid adult animals were placed on a plate, allowed to lay eggs for 3 hours, then removed. 51 hours post egg lay, plates were scored for percentage of animals that reached L4 stage. At least 15 animals were scored for each plate. Visual scoring was done with the examiner blinded to genotype.

### Auxin-induced depletion

Synchronized L1 animals were placed on seeded NGM plates with or without 1 mM auxin for 48 hours at 20°C. L4 animals were harvested for imaging or lysed for qPCR analysis.

### Immunoprecipitation-mass spectrometry (IP-MS)

Approximately 10^5^ synchronized adult animals (grown for 68-72 hours from L1 stage at 20°C) were used in each sample. Animals were harvested and washed three times with 10 mL M9 buffer, then flash frozen in liquid nitrogen and stored at -80°C until use. To prepare worm lysates, frozen worm beads were ground into fine powder with liquid nitrogen-cooled mortar and pestle. The worm powder was transferred into 10 mL of lysis buffer (10 mM HEPES-NaOH (pH 7.5), 50 mM NaCl, 2.5 mM MgCl_2_, 1 mM EDTA, 10% Glycerol, 0.5 mM DTT, 1 mM phenylmethylsulfonyl fluoride, 0.25% Triton X-100) supplemented with 1x cOmplete protease inhibitor cocktail (Roche, 11697498001). After rotating at 4°C for 15 minutes, the lysate was briefly centrifuged and then passed through a 0.45 µm syringe filter (Millipore, SLHPR33RS). Lysate was then incubated with either anti-HA agarose beads (Millipore, A2095) or anti-HA magnetic beads (Thermo Scientific, 88836) at 4°C for 3-4 hours and washed four times with 1mL lysis buffer. To elute bound proteins, beads were incubated with 0.5 M NH_4_OH at 37°C for 20 minutes. The eluate was vacuum dried and resuspended in 100 µl 10 mM Tris-HCl (pH 8.5) and sent to The Taplin Biological Mass Spectrometry Facility at Harvard Medical School for analysis.

### RNA immunoprecipitation (RIP)

Approximately 10,000 synchronized L4 animals of each genotype were flash frozen in liquid nitrogen. Worm pellets were ground to powder with liquid nitrogen-cooled mortar and pestle, and resuspended with lysis buffer (20 mM Tris-HCl, pH7.5, 200 mM NaCl, 2.5 mM MgCl_2_, 10% glycerol, and 0.5% IGEPAL CA-630 supplemented with 1x cOmplete protease inhibitor cocktail (Roche, 11697498001) and 80 U/ml RNAseOUT (Invitrogen, 10777019)). The lysate was passed through a 0.45 μm syringe filter (Millipore, SLHPR33RS), then incubated with anti-HA magnetic beads (Thermo Scientific, 88836) at 4°C for 2 hours. After washing three times with 1 ml lysis buffer, 10% of the beads was saved for immunoblotting. TRIzol (Invitrogen, 15596026) was added directly to the rest of the beads for RNA extraction. Primers used for *Tc1* RNA and cdc-42 mRNA detection are listed in Supplementary Table 1.

### RT-qPCR for spliced/total *Tc1* RNA levels and RT-PCR to detect *Tc1* RNA splicing

Synchronized animals were frozen in TRIzol and total RNA was extracted using ZYMO Direct-zol RNA extraction kit with on-column DNase I digestion. cDNA synthesis was performed using SuperScript™ III First-Strand Synthesis System (Invitrogen, 18080051) with random hexamers following manufacturer’s protocols. Threshold cycle (Ct) numbers were obtained using iTaq Universal SYBR Green Supermix (Bio-Rad, 1725120) on Bio-Rad CFX Duet Real-Time PCR system (Bio-Rad, 12016265). For RT-PCR to check intron splicing efficiency, PCR products were resolved on 2-4% agarose gels. Primers used are listed in Supplementary Table 1.

## Author Contributions

Conceptualization: C.C., S.K,; Investigation: C.C., D.C., D.J.P; Methodology: C.C., D.C.; Formal analysis: C.C., D.C.; Writing: C.C., S.K.; Supervision: S.K.; Funding acquisition: S.K.

## Data Availability

Strains and plasmids are available upon request. Strains used in this study are listed and described in Supplementary Table 2. Complete data of HRPC-1::3xHA IP-MS (two replicates) is available in Supplementary Table 1.

## Acknowledgements

We thank past and previous members of the Kennedy lab for helpful discussions and comments. We thank Dr. David Lowe and Dr. Gang Wan for help on mutant mapping genome analysis. We thank the BPF Genomics Core Facility at Harvard Medical School for their expertise and instrument availability that supported this work.

## Supplemental Information

Figures S1-S9. Supplementary Tables 1-2.

## References

1. Huang CRL, Burns KH, Boeke JD. Active transposition in genomes. Annu Rev Genet. 2012;46:651–75.

2. Klein SJ, O’Neill RJ. Transposable elements: genome innovation, chromosome diversity, and centromere conflict. Chromosome Res. 2018;26:5–23.

3. Slotkin RK, Martienssen R. Transposable elements and the epigenetic regulation of the genome. Nat Rev Genet. 2007;8:272–85.

4. Levin HL, Moran JV. Dynamic interactions between transposable elements and their hosts. Nat Rev Genet. 2011;12:615–27.

5. Wolf G, Greenberg D, Macfarlan TS. Spotting the enemy within: Targeted silencing of foreign DNA in mammalian genomes by the Krüppel-associated box zinc finger protein family. Mob DNA. 2015;6:17.

6. Deniz Ö, Frost JM, Branco MR. Regulation of transposable elements by DNA modifications. Nat Rev Genet. 2019;20:417–31.

7. Cordaux R, Batzer MA. The impact of retrotransposons on human genome evolution. Nat Rev Genet. 2009;10:691–703.

8. Feschotte C, Pritham EJ. DNA transposons and the evolution of eukaryotic genomes. Annu Rev Genet. 2007;41:331–68.

9. Fischer SEJ. Activity and Silencing of Transposable Elements in C. elegans. DNA. 2024;4:129–40.

10. Plasterk RH. The Tc1/mariner transposon family. Curr Top Microbiol Immunol. 1996;204:125–43.

11. Fischer SEJ, Wienholds E, Plasterk RHA. Continuous exchange of sequence information between dispersed Tc1 transposons in the Caenorhabditis elegans genome. Genetics. 2003;164:127–34.

12. Haig D. Transposable elements: Self-seekers of the germline, team-players of the soma. Bioessays. 2016;38:1158–66.

13. Lee E, Iskow R, Yang L, Gokcumen O, Haseley P, Luquette LJ 3rd, et al. Landscape of somatic retrotransposition in human cancers. Science. 2012;337:967–71.

14. Scott EC, Gardner EJ, Masood A, Chuang NT, Vertino PM, Devine SE. A hot L1 retrotransposon evades somatic repression and initiates human colorectal cancer. Genome Res. 2016;26:745–55.

15. Rodić N, Sharma R, Sharma R, Zampella J, Dai L, Taylor MS, et al. Long interspersed element-1 protein expression is a hallmark of many human cancers. Am J Pathol. 2014;184:1280–6.

16. Payer LM, Burns KH. Transposable elements in human genetic disease. Nat Rev Genet. 2019;20:760–72.

17. Teixeira FK, Okuniewska M, Malone CD, Coux R-X, Rio DC, Lehmann R. piRNA-mediated regulation of transposon alternative splicing in the soma and germ line [Internet]. Nature. 2017. p. 268–72. Available from: 10.1038/nature25018

18. Pélisson A, Song SU, Prud’homme N, Smith PA, Bucheton A, Corces VG. Gypsy transposition correlates with the production of a retroviral envelope-like protein under the tissue-specific control of the Drosophila flamenco gene. EMBO J. 1994;13:4401–11.

19. Eide D, Anderson P. Insertion and excision of Caenorhabditis elegans transposable element Tc1. Mol Cell Biol. 1988;8:737–46.

20. Eide D, Anderson P. Transposition of Tc1 in the nematode Caenorhabditis elegans. Proc Natl Acad Sci U S A. 1985;82:1756–60.

21. Collins J, Saari B, Anderson P. Activation of a transposable element in the germ line but not the soma of Caenorhabditis elegans. Nature. 1987;328:726–8.

22. Beyer AL, Christensen ME, Walker BW, LeStourgeon WM. Identification and characterization of the packaging proteins of core 40S hnRNP particles. Cell. 1977;11:127–38.

23. König J, Zarnack K, Rot G, Curk T, Kayikci M, Zupan B, et al. iCLIP reveals the function of hnRNP particles in splicing at individual nucleotide resolution. Nat Struct Mol Biol. 2010;17:909–15.

24. Fischl H, Neve J, Wang Z, Patel R, Louey A, Tian B, et al. hnRNPC regulates cancer-specific alternative cleavage and polyadenylation profiles. Nucleic Acids Res. 2019;47:7580–91.

25. Gruber AJ, Schmidt R, Gruber AR, Martin G, Ghosh S, Belmadani M, et al. A comprehensive analysis of 3’ end sequencing data sets reveals novel polyadenylation signals and the repressive role of heterogeneous ribonucleoprotein C on cleavage and polyadenylation. Genome Res. 2016;26:1145–59.

26. Deininger P. Alu elements: know the SINEs. Genome Biol. 2011;12:236.

27. Zarnack K, König J, Tajnik M, Martincorena I, Eustermann S, Stévant I, et al. Direct competition between hnRNP C and U2AF65 protects the transcriptome from the exonization of Alu elements. Cell. 2013;152:453–66.

28. Wilusz J, Shenk T. A uridylate tract mediates efficient heterogeneous nuclear ribonucleoprotein C protein-RNA cross-linking and functionally substitutes for the downstream element of the polyadenylation signal. Mol Cell Biol. 1990;10:6397–407.

29. Kim JH, Paek KY, Choi K, Kim T-D, Hahm B, Kim K-T, et al. Heterogeneous nuclear ribonucleoprotein C modulates translation of c-myc mRNA in a cell cycle phase-dependent manner. Mol Cell Biol. 2003;23:708–20.

30. Shetty S. Regulation of urokinase receptor mRNA stability by hnRNP C in lung epithelial cells. Mol Cell Biochem. 2005;272:107–18.

31. Ray D, Kazan H, Cook KB, Weirauch MT, Najafabadi HS, Li X, et al. A compendium of RNA-binding motifs for decoding gene regulation. Nature. 2013;499:172–7.

32. Cieniková Z, Damberger FF, Hall J, Allain FH-T, Maris C. Structural and mechanistic insights into poly(uridine) tract recognition by the hnRNP C RNA recognition motif. J Am Chem Soc. 2014;136:14536–44.

33. Swanson MS, Dreyfuss G. Classification and purification of proteins of heterogeneous nuclear ribonucleoprotein particles by RNA-binding specificities. Mol Cell Biol. 1988;8:2237–41.

34. Harris LJ, Rose AM. Somatic excision of transposable element Tc1 from the Bristol genome of Caenorhabditis elegans. Mol Cell Biol. 1986;6:1782–6.

35. Eide D, Anderson P. Transposition of Tc1 in the nematode Caenorhabditis elegans. Proc Natl Acad Sci U S A. 1985;82:1756–60.

36. Collins J, Saari B, Anderson P. Activation of a transposable element in the germ line but not the soma of Caenorhabditis elegans. Nature. 1987;328:726–8.

37. Moerman DG, Waterston RH. Spontaneous unstable unc-22 IV mutations in C. elegans var. Bergerac. Genetics. 1984;108:859–77.

38. Vastenhouw NL, Fischer SEJ, Robert VJP, Thijssen KL, Fraser AG, Kamath RS, et al. A genome-wide screen identifies 27 genes involved in transposon silencing in C. elegans. Curr Biol. 2003;13:1311–6.

39. Preston MA, Porter DF, Chen F, Buter N, Lapointe CP, Keles S, et al. Unbiased screen of RNA tailing activities reveals a poly(UG) polymerase. Nat Methods. 2019;16:437–45.

40. Shukla A, Yan J, Pagano DJ, Dodson AE, Fei Y, Gorham J, et al. poly(UG)-tailed RNAs in genome protection and epigenetic inheritance. Nature. 2020;582:283–8.

41. Sijen T, Plasterk RHA. Transposon silencing in the Caenorhabditis elegans germ line by natural RNAi. Nature. 2003;426:310–4.

42. Chen C-CG, Simard MJ, Tabara H, Brownell DR, McCollough JA, Mello CC. A member of the polymerase beta nucleotidyltransferase superfamily is required for RNA interference in C. elegans. Curr Biol. 2005;15:378–83.

43. Zhang C, Montgomery TA, Gabel HW, Fischer SEJ, Phillips CM, Fahlgren N, et al. mut-16 and other mutator class genes modulate 22G and 26G siRNA pathways in Caenorhabditis elegans. Proc Natl Acad Sci U S A. 2011;108:1201–8.

44. Phillips CM, Montgomery TA, Breen PC, Ruvkun G. MUT-16 promotes formation of perinuclear mutator foci required for RNA silencing in the C. elegans germline. Genes Dev. 2012;26:1433–44.

45. Dodson AE, Kennedy S. Phase Separation in Germ Cells and Development. Dev Cell. 2020;55:4–17.

46. Spike CA, Bader J, Reinke V, Strome S. DEPS-1 promotes P-granule assembly and RNA interference in C. elegans germ cells. Development. 2008;135:983–93.

47. Wan G, Fields BD, Spracklin G, Shukla A, Phillips CM, Kennedy S. Spatiotemporal regulation of liquid-like condensates in epigenetic inheritance. Nature. 2018;557:679–83.

48. Dodson AE, Kennedy S. Phase Separation in Germ Cells and Development. Dev Cell. 2020;55:4–17.

49. Guang S, Bochner AF, Pavelec DM, Burkhart KB, Harding S, Lachowiec J, et al. An Argonaute transports siRNAs from the cytoplasm to the nucleus. Science. 2008;321:537–41.

50. Bosher JM, Dufourcq P, Sookhareea S, Labouesse M. RNA Interference Can Target Pre-mRNA: Consequences for Gene Expression in a Caenorhabditis elegans Operon. Genetics. 1999;153:1245–56.

51. Guang S, Bochner AF, Burkhart KB, Burton N, Pavelec DM, Kennedy S. Small regulatory RNAs inhibit RNA polymerase II during the elongation phase of transcription. Nature. 2010;465:1097–101.

52. Vos JC, van Luenen HG, Plasterk RH. Characterization of the Caenorhabditis elegans Tc1 transposase in vivo and in vitro. Genes Dev. 1993;7:1244–53.

53. Sijen T, Plasterk RHA. Transposon silencing in the Caenorhabditis elegans germ line by natural RNAi. Nature. 2003;426:310–4.

54. Fischer SEJ. Activity and Silencing of Transposable Elements in C. elegans. DNA. 2024;4:129–40.

55. Wan G, Yan J, Fei Y, Pagano DJ, Kennedy S. A conserved NRDE-2/MTR-4 complex mediates nuclear RNAi in Caenorhabditis elegans. Genetics. 2020;216:1071–85.

56. Billi AC, Fischer SEJ, Kim JK. Endogenous RNAi pathways in C. elegans. WormBook. 2014;1– 49.

57. McCloskey A, Taniguchi I, Shinmyozu K, Ohno M. hnRNP C tetramer measures RNA length to classify RNA polymerase II transcripts for export. Science. 2012;335:1643–6.

58. Liu N, Dai Q, Zheng G, He C, Parisien M, Pan T. N(6)-methyladenosine-dependent RNA structural switches regulate RNA-protein interactions. Nature. 2015;518:560–4.

59. Laski FA, Rio DC, Rubin GM. Tissue specificity of Drosophila P element transposition is regulated at the level of mRNA splicing. Cell. 1986;44:7–19.

60. Ghanim GE, Rio DC, Teixeira FK. Mechanism and regulation of P element transposition. Open Biol. 2020;10:200244.

61. Tabara H, Sarkissian M, Kelly WG, Fleenor J, Grishok A, Timmons L, et al. The rde-1 gene, RNA interference, and transposon silencing in C. elegans. Cell. 1999;99:123–32.

62. Brenner S. The genetics of Caenorhabditis elegans. Genetics. 1974;77:71–94.

63. Arribere JA, Bell RT, Fu BXH, Artiles KL, Hartman PS, Fire AZ. Efficient marker-free recovery of custom genetic modifications with CRISPR/Cas9 in Caenorhabditis elegans. Genetics. 2014;198:837–46.

64. Ghanta KS, Mello CC. Melting dsDNA donor molecules greatly improves precision genome editing in Caenorhabditis elegans. Genetics. 2020;216:643–50.

65. Concordet J-P, Haeussler M. CRISPOR: intuitive guide selection for CRISPR/Cas9 genome editing experiments and screens. Nucleic Acids Res. 2018;46:W242–5.

66. Schwartz ML, Jorgensen EM. SapTrap, a Toolkit for High-Throughput CRISPR/Cas9 Gene Modification in Caenorhabditis elegans. Genetics. 2016;202:1277–88.

67. Dickinson DJ, Pani AM, Heppert JK, Higgins CD, Goldstein B. Streamlined Genome Engineering with a Self-Excising Drug Selection Cassette. Genetics. 2015;200:1035–49.

68. Frøkjær-Jensen C, Davis MW, Ailion M, Jorgensen EM. Improved Mos1-mediated transgenesis in C. elegans. Nat Methods. 2012;9:117–8.

69. Frøkjær-Jensen C, Davis MW, Sarov M, Taylor J, Flibotte S, LaBella M, et al. Random and targeted transgene insertion in Caenorhabditis elegans using a modified Mos1 transposon. Nat Methods. 2014;11:529–34.

70. Schindelin J, Arganda-Carreras I, Frise E, Kaynig V, Longair M, Pietzsch T, et al. Fiji: an open-source platform for biological-image analysis. Nat Methods. 2012;9:676–82.

71. Eide D, Anderson P. The gene structures of spontaneous mutations affecting a Caenorhabditis elegans myosin heavy chain gene. Genetics. 1985;109:67–79.

72. Livak KJ, Schmittgen TD. Analysis of Relative Gene Expression Data Using Real-Time Quantitative PCR and the 2−ΔΔCT Method [Internet]. Methods. 2001. p. 402–8. Available from: 10.1006/meth.2001.1262

73. Waterhouse AM, Procter JB, Martin DMA, Clamp M, Barton GJ. Jalview Version 2--a multiple sequence alignment editor and analysis workbench. Bioinformatics. 2009;25:1189–91.

74. Troshin PV, Procter JB, Barton GJ. Java bioinformatics analysis web services for multiple sequence alignment--JABAWS:MSA. Bioinformatics. 2011;27:2001–2.

75. Edgar RC. MUSCLE: multiple sequence alignment with high accuracy and high throughput. Nucleic Acids Res. 2004;32:1792–7.

76. UniProt Consortium. UniProt: The universal protein knowledgebase in 2025. Nucleic Acids Res. 2025;53:D609–17.

