## Supplemental Information for "Transposon Regulation in the *Caenorhabditis elegans* Germline and Soma"

**Figures S1-S9.**

**Supplementary Table 1: Complete data of HRPC-1::3xHA IP-MS (two replicates)**

**Supplementary Table 2: List of primers and strains**

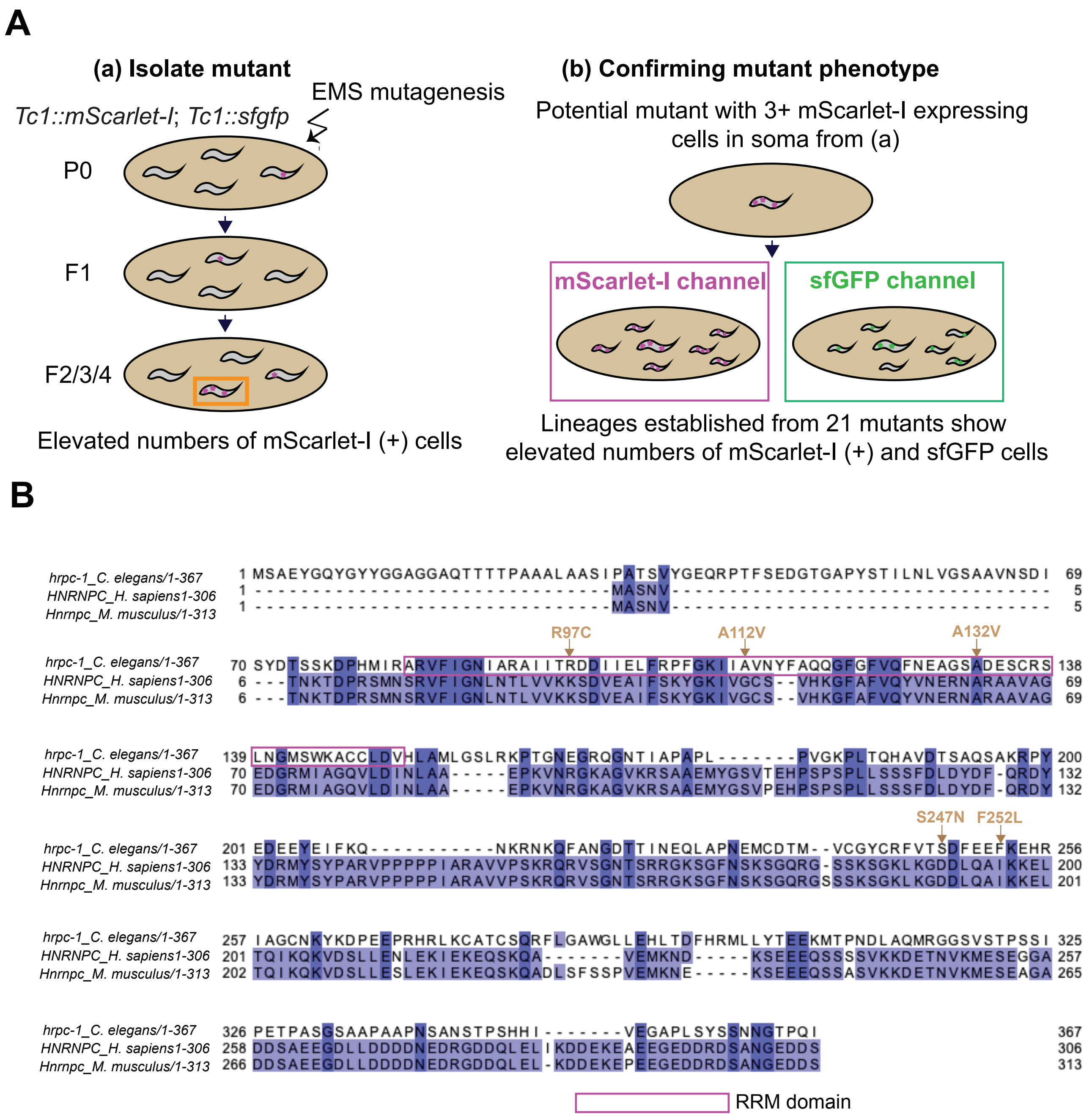


**Figure S1. A genetic screen identifies alleles that increase *Tc1* excision in the soma. Related to Figure 3. (A)** Design of ethyl methanesulfonate (EMS) mutagenesis screen to identify suppressors of somatic *Tc1* excision. Step (a): *Tc1::mScarlet-I*; *Tc1::sfgfp* strains were mutagenized and mutants displaying increased numbers of somatic cells expressing mScarlet-I were isolated at F2, F3 or F4 stages. Step (b): Mutant phenotypes were confirmed by the presence of elevated somatic cells expressing mScarlet-I and sfGFP in progeny of isolated mutants. **(B)** Alignment of *C. elegans* HRPC-1, human HNRNPC, and mouse Hnrnpc. Alleles identified in EMS screens are labeled with brown arrows. RNA recognition motif (RRM) is bracketed in purple.

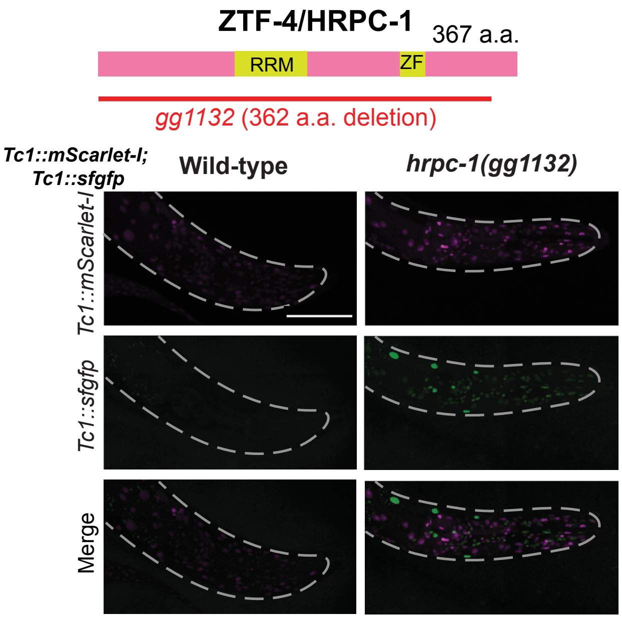


**Figure S2. HRPC-1 deletion allele recapitulates the enhanced soma *Tc1* excision phenotype. Related to Figure 3.** Fluorescent micrographs of *hrpc-1(gg1132)* animals exhibiting increased somatic *Tc1* excision, indicated by the increase in number of somatic cells expressing mScarlet-I and sfGFP. Scale bar, 100 μm. *gg1132* was created by CRISPR-Cas9, which removes amino acids 1-362 of HRPC-1 (3197 base pair deletion).

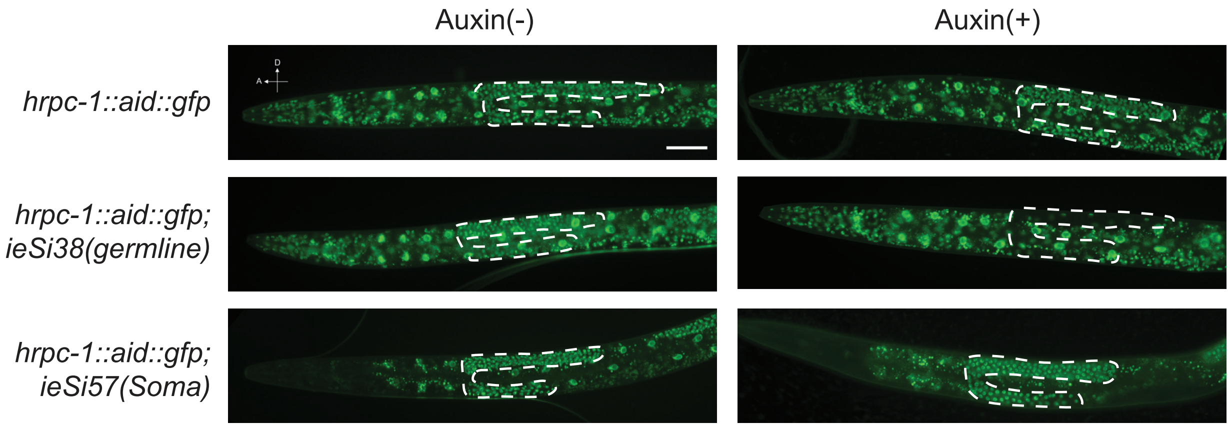


**Figure S3. The auxin-degron system effectively depletes HRPC-1 tissue-specifically. Related to Figure 3.** Representative images for degron-GFP tagged HRPC-1 in strains of indicated genotypes with or without 1 mM auxin treatment. Dotted lines indicate the germline. Note for unknown reasons, a partial diminution of HRPC-1 expression in the soma is observed without auxin treatment in *hrpc-*

*1::aid::gfp; ieSi57*. Animals are at L4 stage. Scale bar, 50 µm. A, anterior; D, dorsal (direction of animal).

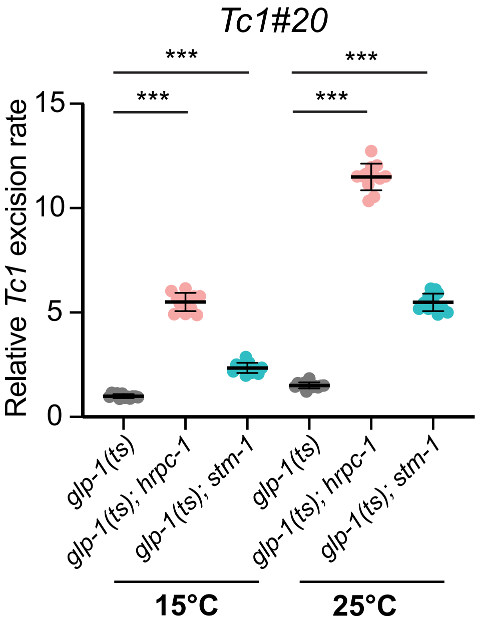


**Figure S4. *Tc1* excision levels are increased in germline-less animals in *hrpc-1* and *stm-1* animals. Related to Figure 3.** qPCR quantification of *Tc1* excision in *glp-1(q224ts)* animals for *Tc1#20* in animals with (15°C) or without (25°C) a germline. For the *glp-1(ts); stm-1* strain*, stm-1* is a 58 base pair deletion in the coding region, which results in a premature *stm-1* stop codon. n=12 per genotype/temperature condition, from 3 biological replicates. ***, p ≤ 0.001 (one-way ANOVA with Dunnett's multiple comparisons test.). Mean ± SD.

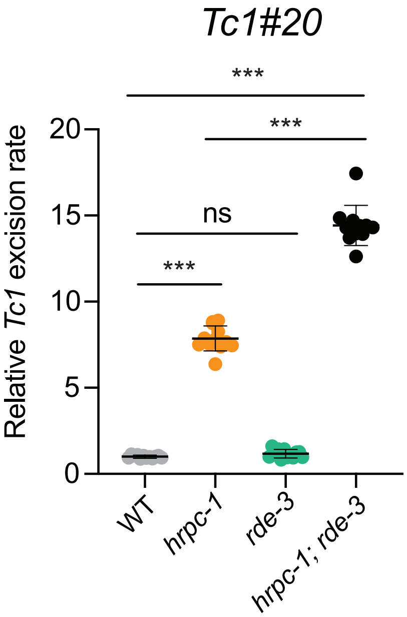

**Figure S5. Increased *Tc1* excision of an endogenous Tc1 element in animals lacking both RNAi and HRPC-1**. **Related to Figure 3.** qPCR quantification of *Tc1* excision for *Tc1#20* in animals of the indicated genotypes. The data shows that *Tc1#20* is excised more in the double mutant than in the single mutants. n=12 per genotype, from 3 biological replicates. ***, p ≤ 0.001 (one-way ANOVA with Dunnett's multiple comparisons test.). Mean ± SD. Note, these results cannot distinguish between somatic or germline excision of *Tc1#20*.

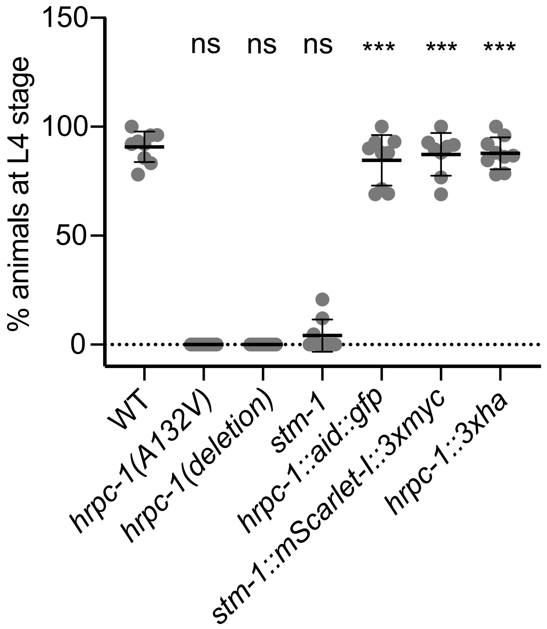


**Figure S6. Epitope tagging does not disrupt HRPC-1 or STM-1 functions. Related to Figure 4 and 5.** Percent of animals reaching the L4 stage within 51-hours after egg-laying at 20°C. *hrpc-1* and *stm-1* mutants show delayed development. Epitope-tagging of *hrpc-1* or *stm-1* does not cause developmental delay. *hrpc-1*(deletion) is *gg1132*. Each data point represents one plate scored; n=8-9 plates scored per genotype, done in 3 biological replicates.

A

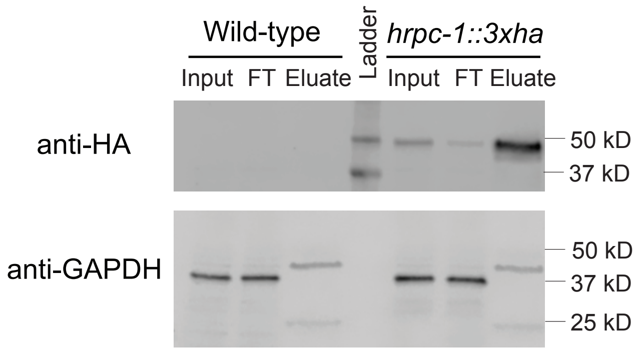


B

| Gene | Control IP (WT) | | *hrpc-1::3xha* IP | |
|---|---|---|---|---|
| | Unique peptides | Total peptides | Unique peptides | Total peptides |
| *stm-1* | 0 | 0 | 21 | 27 |
| *hrpc-1* | 0 | 0 | 9 | 12 |
| *nep-17* | 0 | 0 | 5 | 7 |
| *cpl-1* | 0 | 0 | 5 | 6 |
| *rpl-32* | 0 | 0 | 4 | 4 |

**Figure S7. HRPC-1 interacts with STM-1 *in vivo*. Related to Figure 4.** An independent replicate of the IP-MS experiment shown in Figure 4. **(A)** Gel image of immunoblotting of HA and GAPDH in indicated samples from the IP-MS experiment. Wild-type strain is the control. FT, flow-through fraction. **(B)** Number of unique and total peptides of the top 5 proteins only identified in the *hrpc-1::3xha* IP-MS sample.

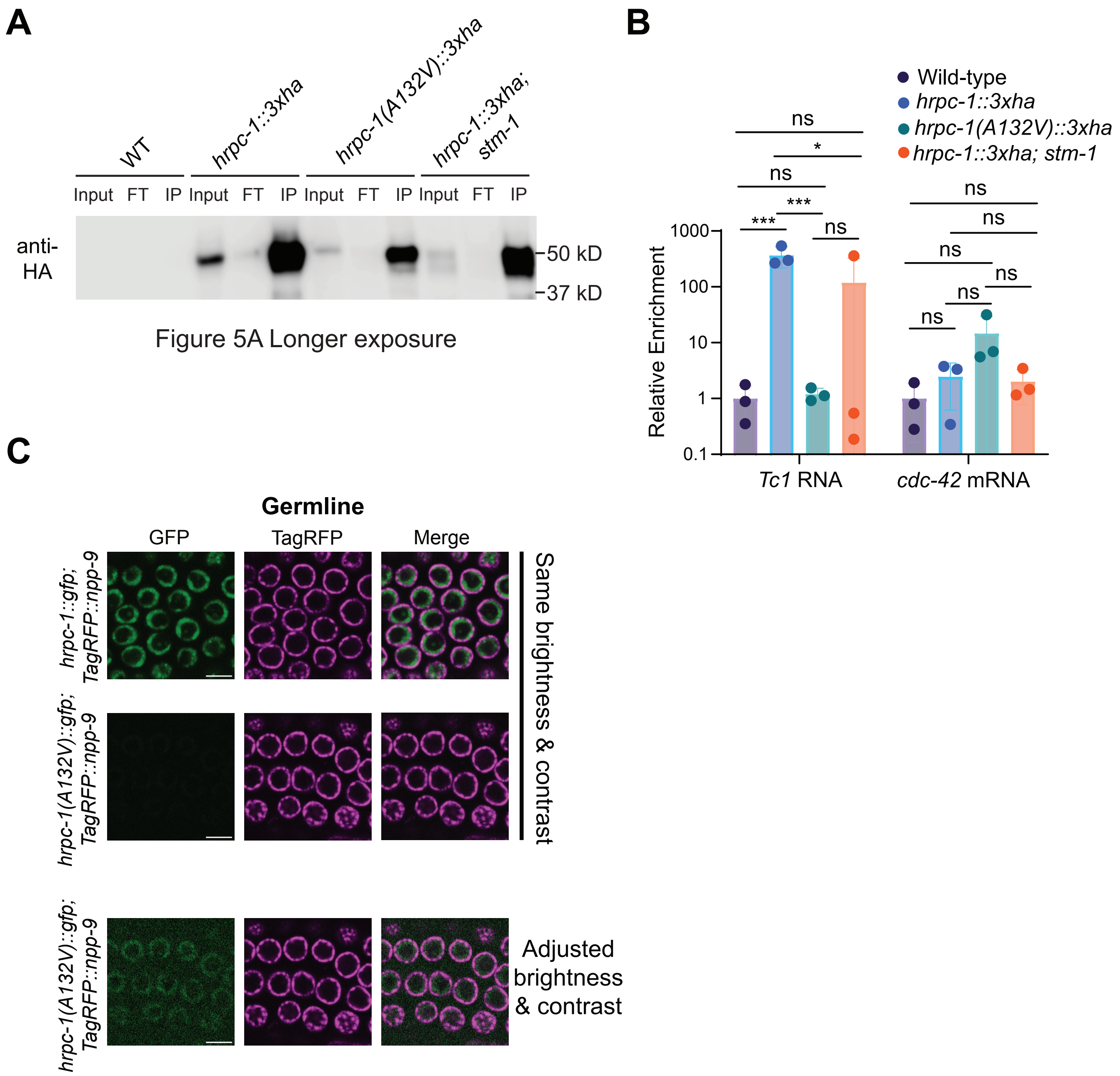


**Figure S8. HRPC-1 binds *Tc1* RNA. Related to Figure 5. (A)** Prolonged exposure of the blot shown in Figure 5A. Note reduced HRPC-1 levels in the input in animals with A132V mutation and *stm-1* mutation. **(B)** Figure 5B results were normalized to the amount of HRPC-1 protein in the IP fraction for corresponding genotypes. Data are from 3 biological replicates. ns: not significant; *, $p \leq 0.05$, ***, $p \leq 0.001$ (Two-way ANOVA with Šídák's multiple comparisons test). Mean ± SD. **(C)** Confocal images showing expression of HRPC-1::GFP and HRPC-1(A132V)::GFP in germ cells. Nuclei are marked with the nuclear pore protein TagRFP::NPP-9. Note the decreased expression of HRPC-1(A132V)::GFP when compared to HRPC-1::GFP, consistent with the decrease in HRPC-1 protein seen in the input lanes of panel **(A)**. Scale bar, 5 µm.

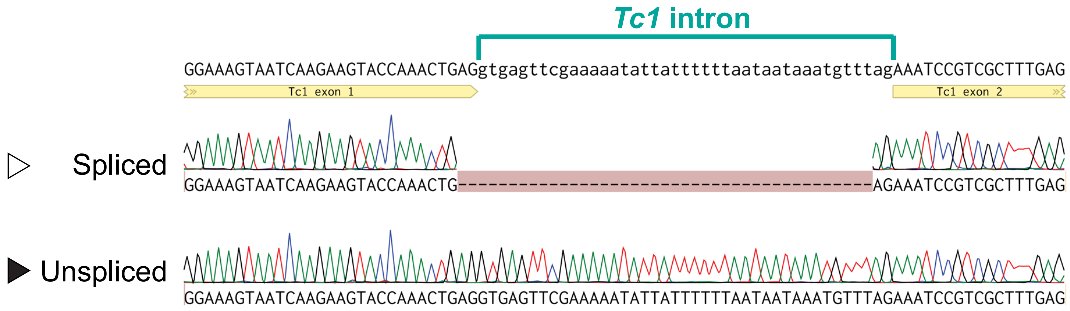


**Figure S9. HRPC-1 and STM-1 affect splicing of *Tc1* RNAs. Related to Figure 6.** Sanger sequencing of RT-PCR products from PCR reactions shown in Figure 6. The sequencing confirms that the top band in Figure 6 is unspliced *Tc1* RNA, and the lower band is spliced *Tc1* RNA.
